# Flexible 3-D Electrochemical Impedance Spectroscopy Sensors Incorporating Phase Delay for Comprehensive Characterization of Atherosclerosis

**DOI:** 10.1101/2023.09.20.558681

**Authors:** Michael Chen, Natalia Neverova, Shili Xu, Krit Suwannaphoom, Gentian Lluri, Mikayla Tamboline, Sandra Duarte, Michael C. Fishbein, Yuan Luo, René R. Sevag Packard

**Author notes:** Correspondence: Dr. René R. Sevag Packard, Departments of Medicine, Physiology, and Bioengineering, UCLA. 10833 Le Conte Ave., CHS Building Room 43-268, Los Angeles, CA 90095.

## Abstract

**Background:** Distinguishing quiescent from rupture-prone atherosclerotic lesions has significant translational and clinical implications. Electrochemical impedance spectroscopy (EIS) characterizes biological tissues by assessing impedance and phase delay responses to alternating current at multiple frequencies.

We evaluated invasive 6-point stretchable EIS sensors over a spectrum of experimental atherosclerosis and compared results with intravascular ultrasound (IVUS), molecular positron emission tomography (PET) imaging, and histology.

**Methods:** Male New Zealand White rabbits (n=16) were placed on a high-fat diet for 4 or 8 weeks, with or without endothelial denudation via balloon injury of the infrarenal abdominal aorta. Rabbits underwent *in vivo* micro-PET imaging of the abdominal aorta with ^68^Ga-DOTATATE, ^18^F-NaF, and ^18^F-FDG, followed by invasive interrogation via IVUS and EIS. Background signal corrected values of impedance and phase delay were determined. Abdominal aortic samples were collected for histological analyses. Analyses were performed blindly.

**Results:** Phase delay correlated with anatomic markers of plaque burden, namely intima/media ratio (r=0.883 at 1 kHz, *P*=0.004) and %stenosis (r=0.901 at 0.25 kHz, *P*=0.002), similar to IVUS. Moreover, impedance was associated with markers of plaque activity including macrophage infiltration (r=0.813 at 10 kHz, *P*=0.008) and macrophage/smooth muscle cell (SMC) ratio (r=0.813 at 25 kHz, *P*=0.026). ^68^Ga-DOTATATE correlated with intimal macrophage infiltration (r=0.861, *P*=0.003) and macrophage/SMC ratio (r=0.831, *P*=0.021), ^18^F-NaF with SMC infiltration (r=-0.842, *P*=0.018), and ^18^F-FDG correlated with macrophage/SMC ratio (r=0.787, *P*=0.036).

**Conclusions:** EIS with phase delay integrates key atherosclerosis features that otherwise require multiple complementary invasive and non-invasive imaging approaches to capture. These findings indicate the potential of invasive EIS as a comprehensive modality for evaluation of human coronary artery disease.

**GRAPHICAL ABSTRACT:** 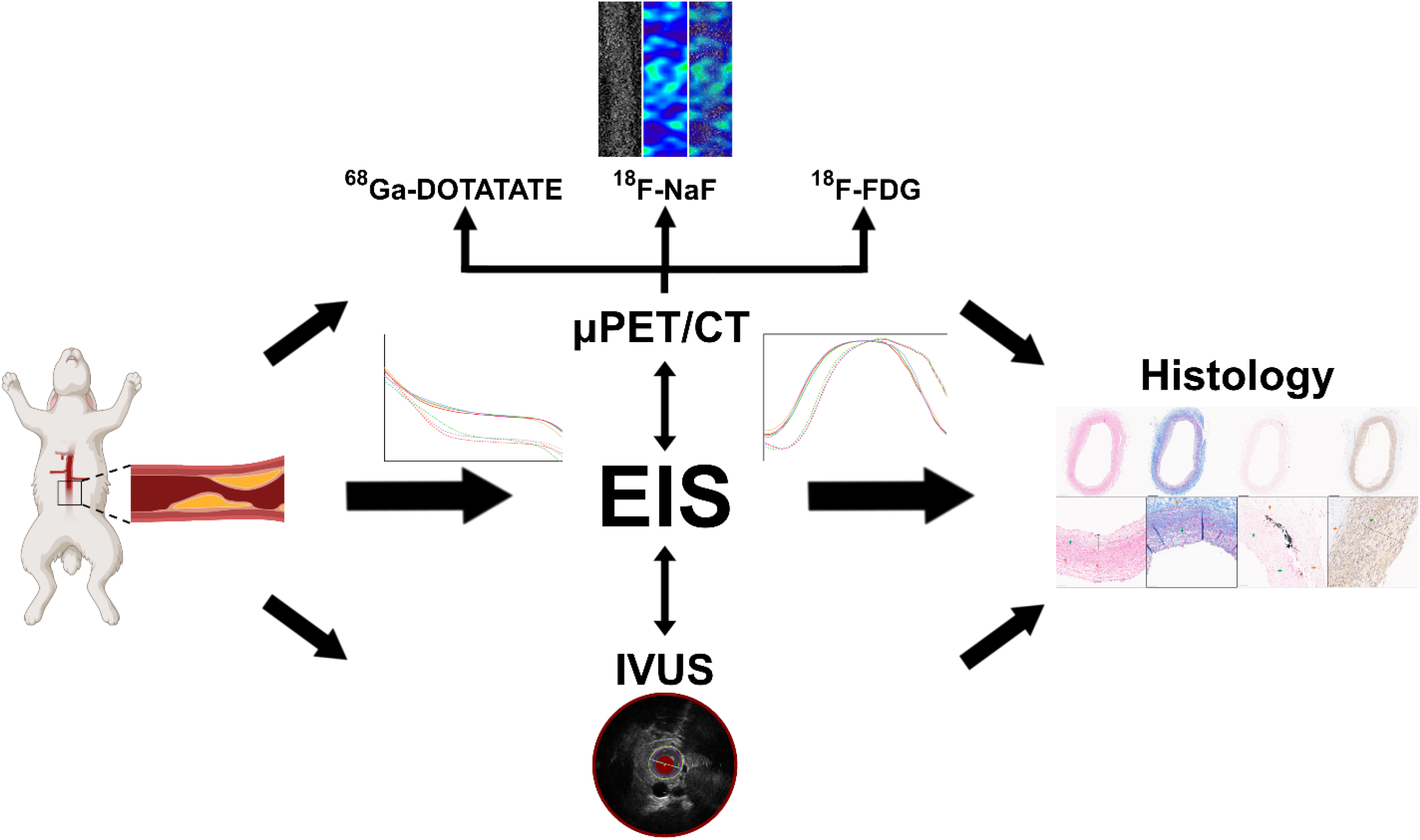

**HIGHLIGHTS:**

- Electrochemical impedance spectroscopy (EIS) characterizes both anatomic features – via phase delay; and inflammatory activity – via impedance profiles, of underlying atherosclerosis.
- EIS can serve as an integrated, comprehensive metric for atherosclerosis evaluation by capturing morphological and compositional plaque characteristics that otherwise require multiple imaging modalities to obtain.
- Translation of these findings from animal models to human coronary artery disease may provide an additional strategy to help guide clinical management.

## INTRODUCTION

Cardiovascular disease remains the leading cause of mortality in the U.S., accounting for 20.1% of all deaths in 2021, often due to coronary artery disease (CAD) complications.^1^ We have significantly progressed in our understanding of the pathobiology of atherosclerosis. In particular, lesional macrophage infiltration, neo-intimal macrophage to vascular smooth muscle cell (SMC) ratio, presence of a necrotic core overlain by a thin fibrous cap, positive remodeling, and spotty calcification, amongst others, are now appreciated as indicators of rupture-prone, inflammatorily active, ‘vulnerable’ plaques.^2^ In contemporary clinical practice promoting aggressive lipid-lowering strategies, we have furthermore witnessed an increase in the incidence of endothelial erosion as opposed to fibrous cap rupture as the underlying mechanism leading to acute coronary syndrome (ACS).^3^ Accordingly, there is significant translational and clinical interest in the development of invasive and non-invasive imaging strategies to reveal pertinent components of atherosclerotic lesions prior to downstream clinical events.

Non-invasive imaging modalities such as coronary computed tomography angiography (CTA) provide comprehensive visualization of the entire coronary tree and the severity, type, calcification, and extent of atherosclerotic plaques, an assessment that can be augmented by fractional flow reserve by computed tomography (FFR-CT).^4^ Positron emission tomography (PET) evaluation of the coronary arteries, beyond myocardial perfusion imaging and myocardial blood flow quantitation^5^, can provide information on specific plaque components such as macrophage infiltration via ^68^Ga-tetraazacyclododecanetetraacetic acid-DPhe1-Tyr3-octreotate (^68^Ga-DOTATATE)^6^, microcalcification via ^18^F-sodium fluoride (^18^F-NaF)^7^, and metabolic/inflammatory activity via ^18^F-fluorodeoxyglucose (^18^F-FDG).^8^ However, non-invasive imaging modalities suffer from drawbacks such as blooming artifact and need for contrast agents in coronary CTA, and the limited spatial resolution and reliance on radiotracers for PET imaging.^9^

Invasive imaging modalities can help guide CAD management by directly characterizing target atherosclerotic lesions. Intravascular ultrasound (IVUS) is a well-established, catheter-based imaging method capable of measuring vessel wall dimensions, determining plaque phenotype such as degree of atherosclerosis burden and calcification, and assessing the distribution and severity of plaques along a vessel. However, IVUS cannot accurately discriminate amongst plaque components and has relatively lower spatial resolution compared to other invasive imaging modalities.^10^ Optical coherence tomography (OCT) complements the strengths and limitations of IVUS. OCT utilizes near-infrared light to generate cross-sectional images of a sample by calculating the delay times between different light rays detected by the sensor, and boasts a 10-fold higher spatial resolution than IVUS, although at the expense of a lower penetration depth.^11,12^ Near-infrared fluorescence (NIRF) also utilizes near-infrared light, which, instead of directly interrogating the sample, stimulates fluorophores that localize and interact with the target of interest. NIRF provides deep tissue penetration, mainly due to low signal attenuation from blood, and lower background noise from tissue autofluorescence, and is capable of assessing plaque inflammation *in vivo.*^13^

Ideally, invasive coronary imaging should elucidate the structural and compositional features of the interrogated atherosclerotic lesion. Electrochemical impedance spectroscopy (EIS) assesses the resistive and charge-storing characteristics of biological tissue by measuring the impedance that develops in response to an applied alternating current (AC). By interrogating a sample over a range of AC frequencies, the frequency-dependent electrical properties of a tissue sample, such as impedance, a measure of resistance to current flow, can be determined.^14^ Our group previously demonstrated that EIS distinguishes lipid-laden atherosclerotic lesions plaques from healthy arterial segments.^15^ Further improvements to the initial linear 2-point sensor yielded a 6-point circumferential design capable of 360° interrogation of the endoluminal surface, thus accounting for the eccentric nature of atherosclerosis.^16^ In the present study, we incorporated background signal correction for impedance, integrated the EIS phase delay, i.e., the offset between input and output signals, and determined the diagnostic performance of invasive EIS over a wide range of atherosclerosis disease severity conditions in a New Zealand White (NZW) rabbit model. We further interrogated resultant plaque phenotypes via ^68^Ga-DOTATATE, ^18^F-NaF, and ^18^F-FDG micro-PET/CT imaging and by IVUS, and conducted histological analyses of pertinent plaque parameters. Our results establish invasive EIS with phase delay as a strategy providing complementary structural and phenotypic atherosclerotic plaque characterization, thereby permitting the distinction between apparently stable and more advanced lesions.

## RESULTS

### Blood Work

Serum C-reactive protein (CRP) and lipid levels at baseline were similar in all animals (**Figure S1, Table S1**). At harvesting, all groups had significantly increased total- and LDL-cholesterol levels. Triglyceride levels increased significantly in the 4-week high-fat & balloon injury (4WK HF/BI) group only, whereas CRP levels did not increase significantly. Intergroup analyses revealed that at the time of harvesting, HDL-cholesterol and triglycerides were lower in the 8-week high-fat & balloon injury (8WK HF/BI) group compared to the 4WK HF/BI group (**Figure S1, Table S1**).

### Histology

Other than the 4-week high-fat (4WK HF) group and one 8-week high-fat (8WK HF) rabbit, intimal thickening was present in the abdominal aortas of all rabbits (**Figures S2, S3**). Whereas the 8WK HF group displayed minute, focal intimal thickening (27.6 ± 6.7 μm), substantial, circumferential intimal thickening was present in both the 4WK HF/BI (125.5 ± 67.8 μm) and 8WK HF/BI (200.6 ± 81.2 μm) groups. Similarly, intima/media ratio and %stenosis, absent in the 4WK HF group, were lowest in the 8WK HF group (0.14 ± 0.03, 1.76% ± 1.76%), followed by the 4WK HF/BI group (0.64 ± 0.51, 4.80% ± 2.07%), and most significant in the 8WK HF/BI group (1.08 ± 0.05, 7.94% ± 2.05%) (**Figure S3A-B**). Medial calcification was detected in n=3 animals: in the 4WK HF/BI group (n=2, 6.41 ± 7.17 x 10^4^ μm^2^) and the 8WK HF/BI group (n=1, 5.98 x 10^4^ μm^2^). In the HF only groups, there was no macrophage or intimal SMC infiltration at 4WK, and these were present only in a subset at 8WK (n=2 for macrophage, 35.00% ± 21.21%). On the other hand, all rabbits from the 4WK HF/BI and 8WK HF/BI groups exhibited macrophage (37.50% ± 21.02%, and 28.33% ± 2.89%, respectively) and SMC (63.61% ± 4.15%, and 55.60% ± 7.56%, respectively) accumulation in the intimal layer (**Figure S3C-E**).

### Micro-PET/CT

Rabbits underwent *in vivo* micro-PET/CT imaging on consecutive days to evaluate macrophage presence by ^68^Ga-DOTATATE, microcalcification by ^18^F-NaF, and metabolic activity by ^18^F-FDG, respectively, within the infrarenal abdominal aortic area of interest (**Figure 3A-B, Figure S4**). The mean % injected dose per cubic centimeter (%ID/cc) of ^68^Ga-DOTATATE strongly correlated with intimal macrophage infiltration (r=0.861, *P*=0.003) and macrophage/SMC ratio (r=0.831, *P*=0.021) (**Figure 3C-D**). Mean %ID/cc TBR of ^18^F-FDG moderately correlated with macrophage/SMC ratio (r=0.787, *P*=0.036) but not with macrophage infiltration alone (r=0.524, *P*=0.147) (**Figure 3E-F**), intima/media ratio (r=-0.311, *P*=0.415), nor %stenosis (r=-0.284, *P*=0.458). Mean %ID/cc TBR of ^18^F-NaF strongly correlated with SMC infiltration (r=-0.842, P=0.018) (**Figure S5**).

## IVUS

Rabbits from the 4WK HF group displayed no luminal narrowing, while those from the 8WK HF/BI group exhibited the highest degree of plaque burden (28.8% ± 1.3%) (**Figure S6**). Comparisons between timepoints (4 vs. 8 weeks) as well as experimental conditions (high-fat only vs. high-fat + balloon injury) yielded statistically significant differences for all comparisons. Both 4WK HF/BI and 8WK HF groups displayed significantly lower plaque burden than the 8WK HF/BI group (*P*=0.002 and *P*=0.001, respectively). Plaque burden determined by IVUS correlated significantly with both intima/media ratio (r=0.939, *P*<0.001) and %stenosis (r=0.892, *P*=0.001) (**Figure 4**).

## EIS Impedance and Phase Delay

- **Determination of Stable Impedance ‘Plateau Region’ for EIS Data Analysis** Whereas biological systems demonstrate both capacitive and resistive behavior, the lower and higher impedance frequency regimes are dominated by either capacitive or resistive behavior, respectively, and thus do not provide reliable measurements. To this end, *in vivo* analyses were performed from 40Hz─40 kHz, which constitutes the “plateau region” over which impedance signals are stable, thus allowing for accurate comparisons between conditions (**Figure 5**).
- **EIS Demonstrates Signal Specificity Between Inflated and Deflated Measurements** There were marked differences in impedance and phase delay under balloon inflation vs. deflation *in vivo*, reflecting changes in endoluminal microsensor contact (**Figure 5A, 5C, 5E, 5G**). Under deflation at 1 kHz, the average impedance and phase delay, across all experimental conditions, was 3308.9 ± 1409.7 Ω and −10.3 ± 4.0°, respectively. In contrast, average impedance and phase delay under inflation at 1 kHz across conditions was 17370.2 ± 6978.4 Ω and −5.0 ± 1.3°, respectively (**Figure S7**). Individually, all experimental groups displayed significant disparities between inflated and deflated impedance (*P*<0.001 for all groups) and phase delay (*P*<0.001 for all groups) measurements at 1 kHz. Following impedance TBR measurement, max TBR – selected from n=15 individual values to capture potential heterogeneities in eccentric atherosclerotic lesions – was superior to max background subtraction correction (BSC) and uncorrected (‘raw’) max impedance in the number and strength of correlations with histological plaque features. For phase delay, the minimal (i.e., most negative) BSC – also selected from n=15 individual values – performed better than TBR and uncorrected phase delay.
- **EIS Measurements are Consistent across Different Catheters** The performance of all the catheters utilized for *in vivo* EIS measurements (labeled Catheters A to D) was assessed separately *ex vivo* under control (**Figure 5A-D**) and high-fat conditions (**Figure 5E-H**). Interrogation of the abdominal aorta in a control rabbit (i.e., not fed a high-fat diet) over a range of frequencies indicated similar impedance results at 100 Hz (*P*=0.438), 1 kHz (*P*=0.647), and 10 kHz (*P*=0.376), as well as similar phase delay results (100 Hz: *P*=0.784; 1 kHz: *P*=0.614; 10 kHz: *P*=0.486) (**Figure 5B, 5D**). This was also the case in the abdominal aorta of an 8-week high-fat diet rabbit, for which neither impedance (100 Hz: *P*=0.446; 1 kHz: *P*=0.321; 10 kHz: *P*=0.401) nor phase delay (100 Hz: *P*=0.397; 1 kHz: *P*=0.454; 10 kHz: *P*=0.582) (**Figure 5F, 5H**) displayed significant differences among the various catheters.
- **Impedance and Phase Delay Characterize Atherosclerosis Composition and Morphology** The EIS stretchable microelectrodes were affixed on an inflatable balloon, permitting capture of impedance and phase delay results under inflated and deflated conditions and derivation of TBR. In addition, the 2 rings of 3 micro-electrodes permit 360° interrogation of the arterial segment of interest in vertical, oblique, and horizontal axes, for a total of n=15 pair-wise permutations (**Figure 2**). Invasive EIS metrics were evaluated *in vivo* for detection of atherosclerotic structural and compositional features. Phase BSC demonstrated significant correlation with intima/media ratio (0.04–2.5 kHz), which peaked at 1 kHz (r=0.883, *P*=0.004) (**Figure 6A**), and with %stenosis (0.04–1.6 kHz), which was highest at 0.25 kHz (r=0.901, *P*=0.002) (**Figure 6B**). Lesions with higher plaque burden exhibited a higher minimum (less negative) phase vs. less developed lesions or normal arteries. Furthermore, impedance TBR exhibited significant correlation with macrophage infiltration across the frequencies 0.4–25 kHz, which peaked at 10 kHz (r=0.813, *P*=0.008) (**Figure 6C**), and with macrophage/SMC ratio at 25 kHz (r=0.813, *P*=0.026) (**Figure 6D**). Lesions with higher inflammatory burden exhibited an increase in impedance values. Thus, whereas EIS phase delay captures pertinent atherosclerotic morphological features, impedance detects key compositional features (**Figure 7**).

## DISCUSSION

An imaging modality capable of detecting prominent upstream ACS features would prove to be a powerful tool for lesion characterization and potential downstream changes in therapy. We previously demonstrated the feasibility of a 2-point sensor for endoluminal interrogation by electroplating of the micro-electrodes with platinum black, thereby greatly reducing contact impedance, particularly at higher frequencies.^15^ This strategy further permits an augmentation of the number of micro-electrodes placed on the balloon, laying the foundation for 3-D interrogation^16^ and detection of eccentric atherosclerotic lesions. In the present body of work, we demonstrate the potential for 3-D EIS to serve as a comprehensive modality for atherosclerotic characterization by illustrating the ability of EIS to (i) determine atherosclerosis structural features including %stenosis and intima/media ratio via phase delay, and (ii) unravel histological markers of high-risk lesions including macrophage infiltration and macrophage/vascular SMC ratio, via impedance sweep. We further indicate (iii) enhanced EIS signal specificity when conducting background correction measures, and (iv) signal stability between various EIS sensors.

EIS is capable of discerning atherosclerotic tissue by frequency-dependent electrical impedance.^14,17^ Initial studies by Süselbeck et al. utilized a linear array of four electrodes to assess *ex vivo* impedance readings of rabbit^14^ and human^18,19^ arteries. However, the linear, four-electrode design presents several issues. The large overall size of the sensors reduces its spatial specificity, and the linear configuration renders it unsuitable for capturing the heterogeneous and eccentric nature of atherosclerosis. Subsequent work by others introduced a concentric, bipolar electrode configuration that drastically reduced the overall sensor dimensions.^17^ The flexible construction and concentric design allowed for improved contact with the uneven endoluminal topography.^17,20,21^ Another design featured two distinct electrodes forming one sensor pair^15^; the increased separation between the electrodes increased signal-to-noise ratio compared to the concentric design and allowed for deeper tissue penetration of signals, thereby increasing the range of frequencies over which the sensor can distinguish between varying tissue composition (e.g., healthy endoluminal surface vs neointima burdened by lipid-laden foam cells).^15,22^ However, both of these designs still did not address one of the main limitations of the original design proposed by Süselbeck. The focal nature of these designs is inadequate for capturing the heterogeneity of atherosclerosis; these sensors would need to be rotated between measurements to account for plaque eccentricity. Catheter rotation while under balloon inflation carries the risk of iatrogenic plaque disruption or vessel wall injury; on the other hand, deflating the balloon prior to rotation might shift the placement of the sensors, thus affecting the accuracy of atherosclerotic assessment at that vessel segment. The 6-point, circumferential design utilized in the present study allows for uninterrupted 360° interrogation of a vessel segment. Furthermore, the 15 unique measurement permutations begotten by the 6-point design, combined with the reduced electrode dimensions, improves spatial specificity by facilitating more granular measurement of a vessel segment.

To assess the *in vivo* detection by 3-D EIS signals of atherosclerotic processes of interest, we first evaluated specific plaque components by relevant PET radiotracers that have been studied extensively in both preclinical and clinical investigations. Plaque macrophage content, and more particularly an increased macrophage/SMC ratio, is a known histological indicator of downstream atherosclerotic complications.^2,3^ ^18^F-FDG did not correlate with macrophage infiltration alone and only moderately with the macrophage/SMC ratio. ^18^F-FDG lacks specificity as any cell type that utilizes glucose for metabolism, such as cardiomyocytes, will uptake ^18^F-FDG, leading to limited signal specificity and significant background noise. Previous research indicated a lack of correlation between ^18^F-FDG uptake and CD68 macrophage staining^23^ and unfavorable cardiovascular disease risk profiles.^24^ To address the issue of non-specificity, ^68^Ga-DOTATATE was developed as a somatostatin receptor 2 (SSTR2) agonist. Given proinflammatory macrophages express SSTR2, ^68^Ga-DOTATATE has higher specificity to image plaque macrophages.^25^ In the present study, ^68^Ga-DOTATATE correlated strongly with both macrophage infiltration and the macrophage/SMC ratio. ^18^F-NaF has emerged as a powerful non-invasive method to determine atherosclerotic lesion activity via PET. We observed a significant, strong correlation between ^18^F-NaF and calcified area. Mechanistically, the ^18^F-fluoride ion exchanges with a hydroxyl group in hydroxyapatite crystals; consequently, ^18^F-NaF is capable of visualizing plaque microcalcification, itself correlated with expansion and thus plaques that are deemed ‘unstable’. Despite our results recapitulating ^18^F-NaF detection of microcalcification, this was present in n=3 rabbits only, limiting our ability to assess EIS detection of microcalcification. We also observed a strong negative correlation between ^18^F-NaF and SMC infiltration into the intima, likely due to their known transdifferentiation into an osteochondrogenic-like cell type that contributes to vascular calcification.^26–30^ Indeed, there is a strong correlation between ^18^F-NaF signal, cardiovascular risk factors, and culprit lesions leading to myocardial infarction.^7^

PET imaging, despite its versatility and clinical applicability, suffers from certain drawbacks, primarily its limited spatial resolution and reliance on radiotracers. The spatial resolution of PET is orders of magnitude worse than those of intravascular imaging modalities. Furthermore, accurate PET imaging relies on the radiotracer localizing with the target of interest and minimal background uptake.^9^ An increasingly utilized alternative to invasive coronary angiography (ICA) is coronary CTA^31^, which allows for detailed assessment of various characteristics, such as calcification, plaque volume, or positive remodeling.^32^ In recent years, CTA has been increasingly paired with fractional flow reserve by computed tomography (FFR-CT)^33^ allowing improvement in its specificity^4^ and the serial assessment of CAD burden.^34,35^ However, CTA exposes patients to ionizing radiation, suffers from blooming artifacts, and is dependent on iodinated contrast agents. Magnetic resonance imaging (MRI) boasts superior tissue contrast compared to CT^2^ and can delineate pertinent plaque features, such as fibrous cap or necrotic core, particularly in the carotids.^2,36^ Assessment of macrophage content has also been demonstrated via dynamic contrast-enhanced MRI.^37^ However, MRI is more suited for larger, “immobile” arteries such as the carotids, sensitive to cardiac and respiratory motion, costly, and not suitable for patients with claustrophobia or metal implants.

IVUS provides morphological assessment of vessel lesions, such as vessel wall dimensions and luminal narrowing. However, IVUS suffers from low spatial resolution and a limited ability to distinguish plaque components.^10^ Recently, IVUS has been paired with complementary imaging modalities that shore up its shortcomings, such as OCT, near infrared spectroscopy (NIRS), or NIRF. OCT provides excellent spatial resolution, but itself lacks sufficient penetration depth; the external elastic membrane cannot be visualized via OCT, thus precluding assessment of true vessel size and plaque burden.^11,12^ NIRS reliably produces semiquantitative information on plaque lipid content, albeit with low image depth resolution, but cannot visualize the lumen or assess outer vessel wall dimensions or plaque burden. Furthermore, NIRS is of limited utility outside semiquantitative plaque lipid content assessment. NIRF captures *in vivo* pathobiological processes and visualizes pertinent biological activity at the plaque and cellular level. Depending on the probe, NIRF may target inflammatory protease activity^10^, oxidized LDL content^38^, or fibrin deposition.^39^ Due to the nature of NIRF, it is entirely dependent on the administration of targeting probes and thus vulnerable to the limitations inherent to a probe and its biodistribution that may reduce plaque TBR.^10,38^ While complementing multiple imaging modalities into a hybrid system provides a more complete picture of atherosclerotic plaque by addressing the weaknesses of individual imaging modalities, this harmony relies on successful co-registration of the individual data sets.

Multiple invasive strategies have undergone clinical scrutiny for characterization of apparently stable atherosclerotic lesions and the prediction of downstream clinical events. Whereas a recent study indicated a benefit of adopting an invasive IVUS or OCT-guided coronary revascularization strategy in complex lesions^40^, the PROSPECT study indicated a low 18.2% positive predictive value of IVUS in detecting lesions that cause adverse clinical events.^41^ Furthermore, the 5-year outcomes of the FAME trial indicated an inability of invasive fractional flow reserve (FFR) to decrease death or myocardial infarction in a hemodynamic-guided coronary revascularization strategy.^42^ An ideal invasive plaque characterization strategy should distinguish between apparently stable vs. metabolically active lesions and help guide medical and/or interventional strategies. In the present experimental study, we demonstrate the ability of 3-D EIS to unravel key atherosclerotic features that would otherwise require multiple PET radiotracers and complementary IVUS for identification.

Our study has limitations. The atherosclerotic lesions were experimentally induced and thus are not necessarily representative of human lesions. Additionally, EIS was measured at one timepoint (animal euthanasia). Future research should assess the ability of serial EIS-derived measures to scrutinize structural and compositional changes occurring during atherosclerosis progression. Furthermore, we posit our results lay the foundation for invasive EIS interrogation in human atherosclerotic lesions, and the determination of clinically relevant threshold values for impedance and phase delay that may help guide coronary revascularization decision-making.

## CONCLUSIONS

EIS distinguishes inflammatorily active from quiescent lesions by detecting key atherosclerosis features that otherwise require multiple imaging modalities to capture. Thus, EIS may be a valuable tool for comprehensive CAD evaluation given its ability to characterize both plaque anatomy and underlying activity.

## METHODS

### Animals

Male NZW rabbits (n=16), ages 12-16 weeks and weighing 3-3.5 kg upon arrival (Crl:KBL, Charles River), were fed a 5% peanut oil and 1% cholesterol high-fat diet (LabDiet) (**Figure 1**). Animals were randomly divided into four groups (n=4 each): 4-week high-fat (4WK HF), 4-week high-fat & balloon injury (4WK HF/BI), 8-week high-fat (8WK HF), and 8-week high-fat & balloon injury (8WK HF/BI). This study was approved by the UCLA Office of Animal Research.

**Figure 1.**
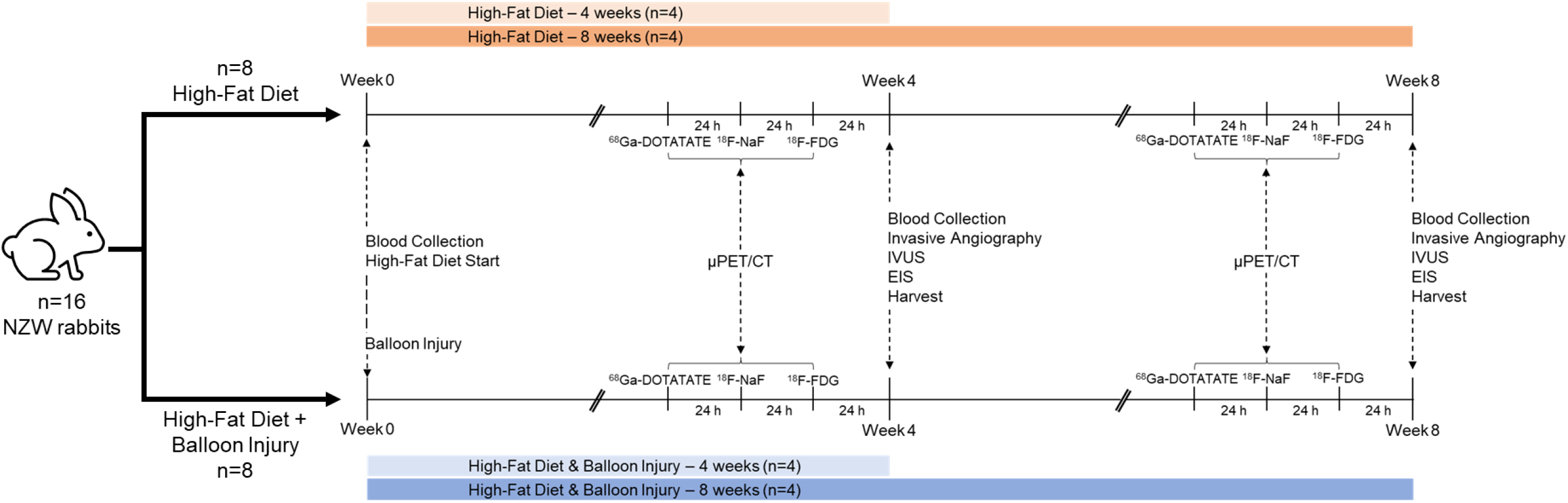
Experimental Timeline. N=16 rabbits were fed a high-fat diet for 4 (n=8) or 8 weeks (n=8), with n=4 from each group undergoing endothelial denudation via balloon injury. PET imaging commenced 3 days prior to harvesting, with each radiotracer (^68^Ga-DOTATATE, ^18^F-NaF, ^18^F-FDG) being imaged 24 h apart. EIS measurements—impedance and phase—were conducted following IVUS. CT: computed tomography. EIS: electrochemical impedance spectroscopy. ^18^F-FDG: ^18^F-fluorodeoxyglucose. ^68^Ga-DOTATATE: ^68^Ga-tetraazacyclododecanetetraacetic acid-DPhe1-Tyr3-octreotate. ^18^F-NaF: ^18^F-sodium fluoride. IVUS: intravascular ultrasound. NZW: New Zealand White. PET: positron emission tomography.

### EIS Sensor Microfabrication

The 6-point EIS sensor was fabricated in-house as previously described.^16^ Briefly, flexible polyimide strips (FPCexpress) with exposed copper pads (600 μm x 300 μm) serving as the electrodes were mounted onto an inflatable balloon (Poba Medical) (15 mm in length, <1 mm diameter under deflation, and ∼4.5 mm under inflation), which was affixed onto the distal end of the catheter tubing (25 cm in length) (Nordson Medical). Tantalum foils (1 mm x 1 mm) (Advent Research Materials) were placed immediately distal and proximal to the balloon to serve as radiopaque markers (**Figure 2**). Insulated copper wires were soldered onto the proximal contact pads of the flexible sensors to be connected to an impedance analyzer (Interface 1010E, Gamry Instruments). Electroplating was performed in a solution of 0.5% w/v PtCl_4_ at −0.6 V for 30 minutes to minimize contact impedance and improve EIS measurement specificity.

**Figure 2.**
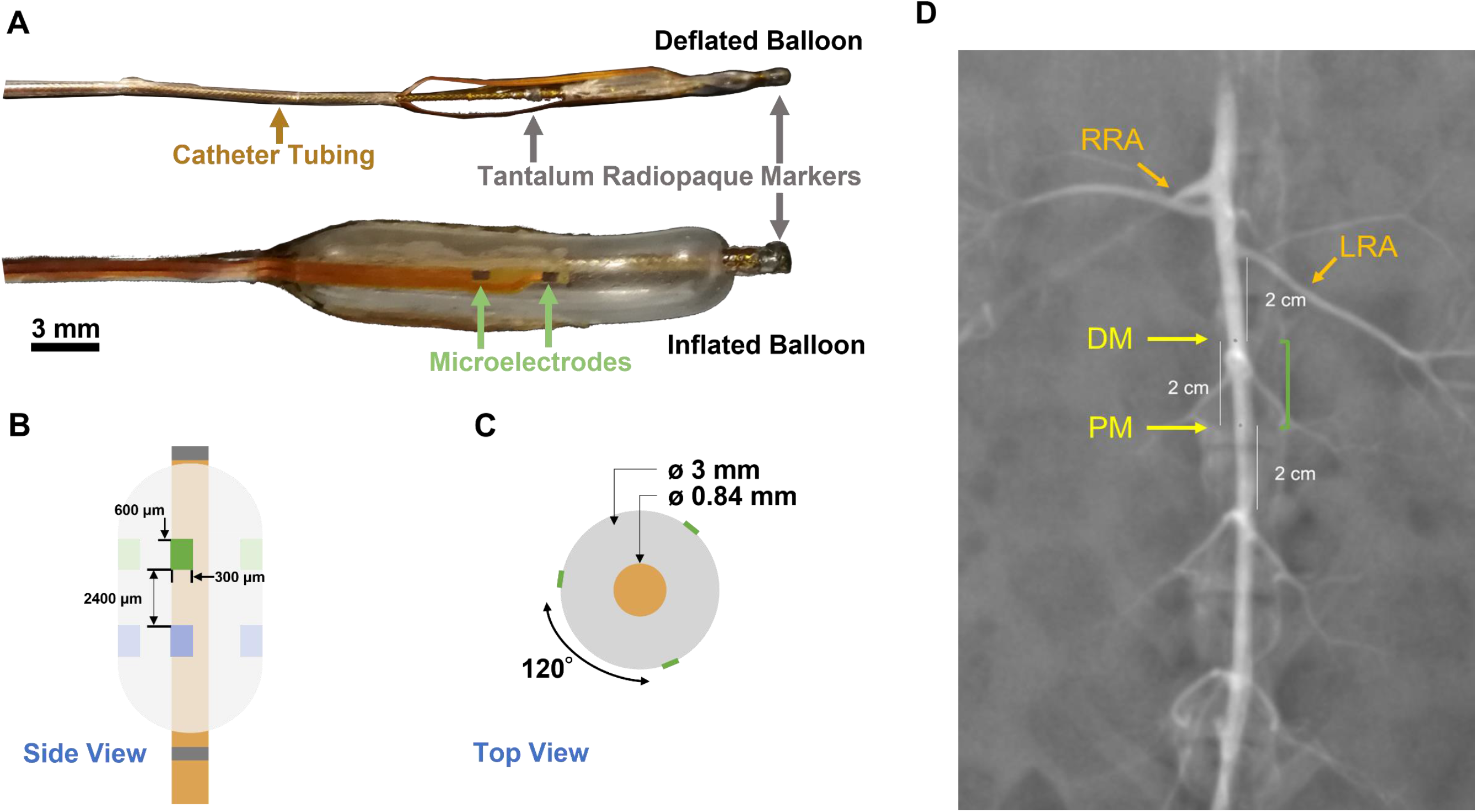
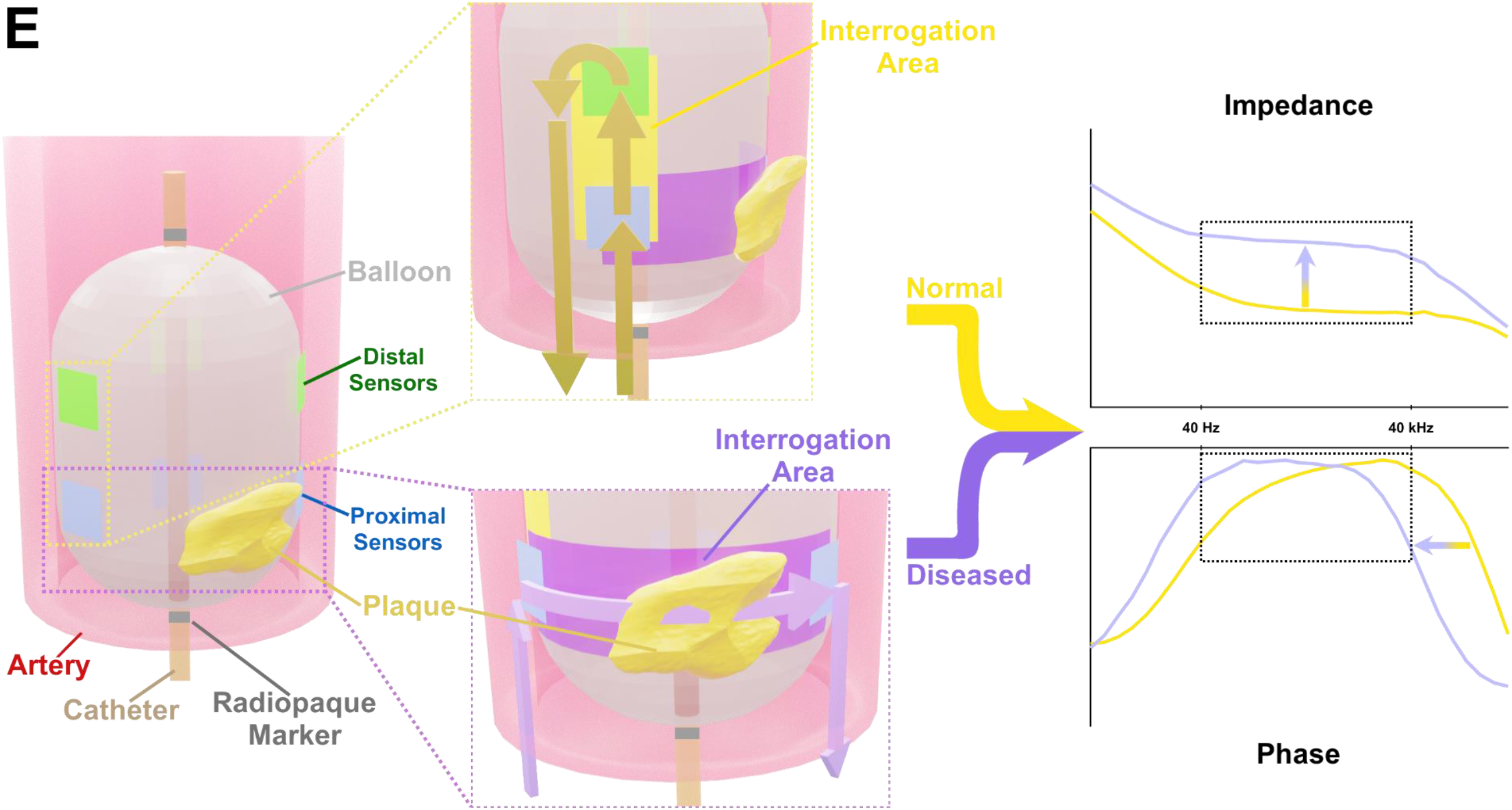
Design and Deployment of the EIS Balloon Catheter. (A) Zoomed-in photos detailing the design of the EIS balloon catheter under deflation and inflation. (B) Side and (C) top-down views of the catheter schematics illustrate the dimensions of the sensors under balloon inflation. (D) Contrast-enhanced invasive angiograms were performed prior to balloon injury, IVUS, and EIS measurements to guide the proper placement of catheters. The image presented displays an angiogram of the infrarenal abdominal aorta prior to EIS measurements. The radiopaque markers (yellow arrows) of the EIS catheter were used to guide the EIS sensors to the proper interrogation site (green backet), consistent with μPET/CT, as measured from the takeoff of the left renal artery (white lines). Guidance of the IVUS catheter was performed in a similar manner. (E) Schematic representation of EIS measurements. The EIS catheter is advanced to the target area under fluoroscopic guidance following which the balloon is inflated for sensor contact with the endoluminal surface. Alternating current (colored 3D arrows) flows through a pair of sensors to generate impedance and phase delay readings for that specific configuration. The 15 unique pairwise sensor configurations produce a comprehensive, 360° interrogation capable of capturing the eccentric nature of atherosclerosis. EIS measurements were performed from 1 Hz to 1 MHz; data from 40 Hz to 40 kHz (‘plateau’ region) was analyzed to assess correlations between impedance or phase delay and plaque characteristics by histology. DM: distal marker. LRA: left renal artery. PM: proximal marker. RRA: right renal artery.

**Figure 3.**
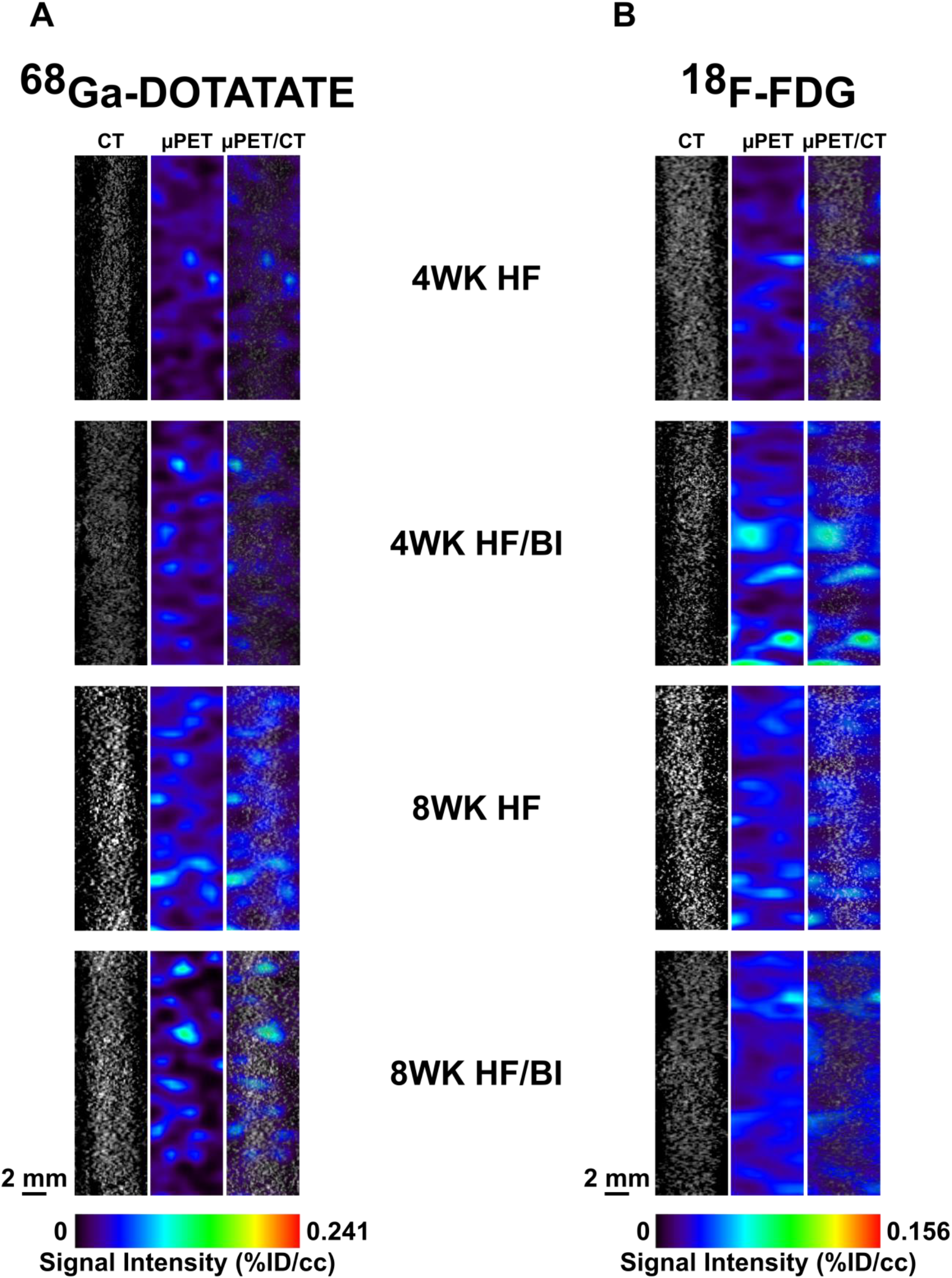
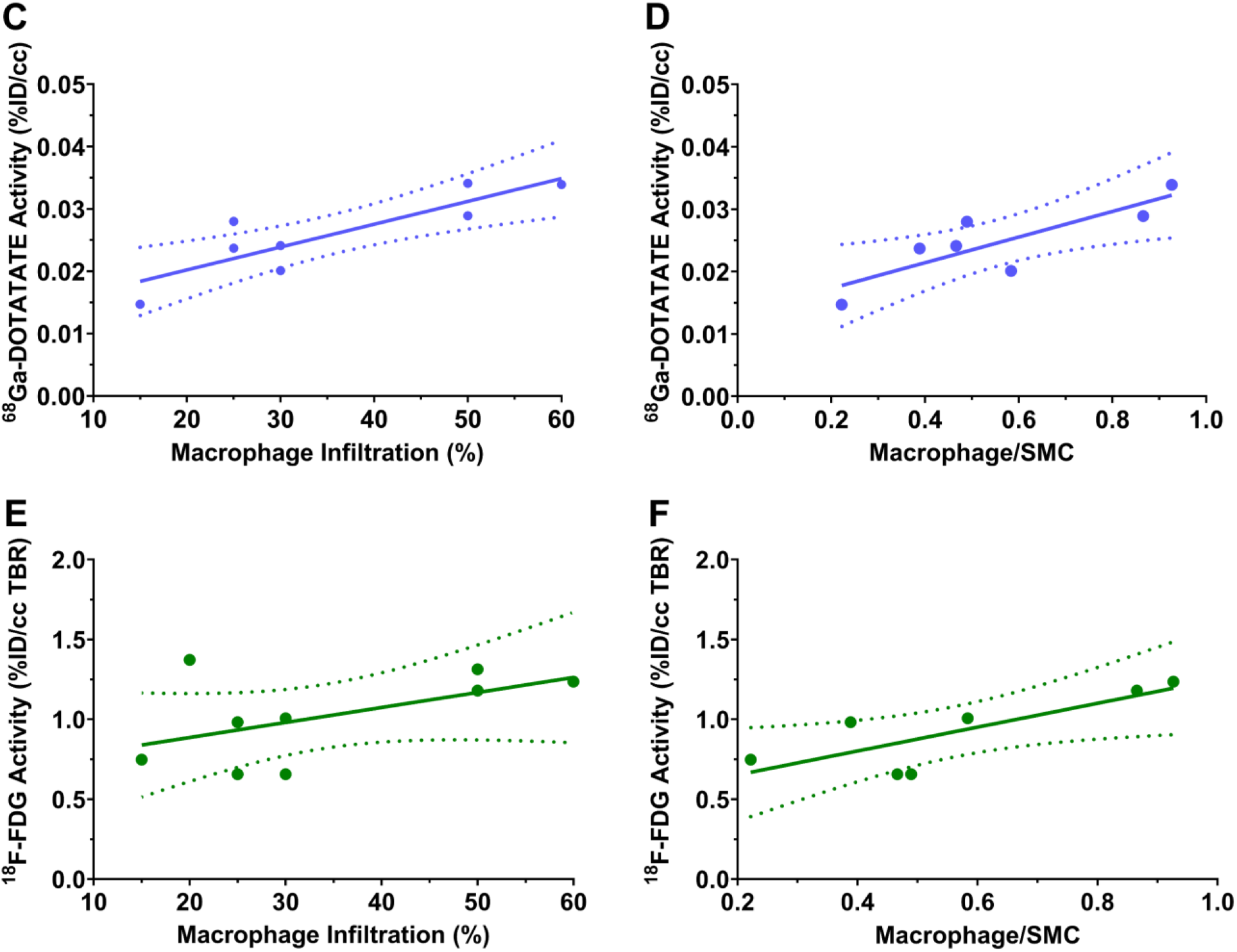
Detection of Atherosclerotic Components by PET Radiotracers. Representative images of the abdominal aorta with (A) ^68^Ga-DOTATATE and (B) ^18^F-FDG obtained on consecutive days to evaluate macrophage presence and metabolic activity. Activities of ^68^Ga-DOTATATE and ^18^F-FDG both correlated with histological plaque parameters of interest. (C, D) Mean %ID/cc of ^68^Ga-DOTATATE strongly correlated with intimal macrophage infiltration (r=0.861, *P*=0.003) and macrophage/SMC ratio (r=0.831, *P*=0.021). (E, F) Mean ^18^F-FDG TBR trended toward correlation with macrophage infiltration (r=0.524, *P*=0.147) without statistical significance, but did moderately correlate with macrophage/SMC ratio (r=0.787, *P*=0.036). Pearson correlation coefficients were calculated for comparison of PET radiotracer activity against histological parameters. For each radiotracer, all CT images and all PET images were obtained using the same scale. ^18^F-FDG: ^18^F-fluorodeoxyglucose. ^68^Ga-DOTATATE: ^68^Ga-tetraazacyclododecanetetraacetic acid-DPhe1-Tyr3-octreotate. HF: high-fat diet. HF/BI: high-fat diet & balloon injury. %ID/cc: % injected dose per cubic centimeter. SMC: smooth muscle cell. TBR: target-to-background ratio.

**Figure 4.**
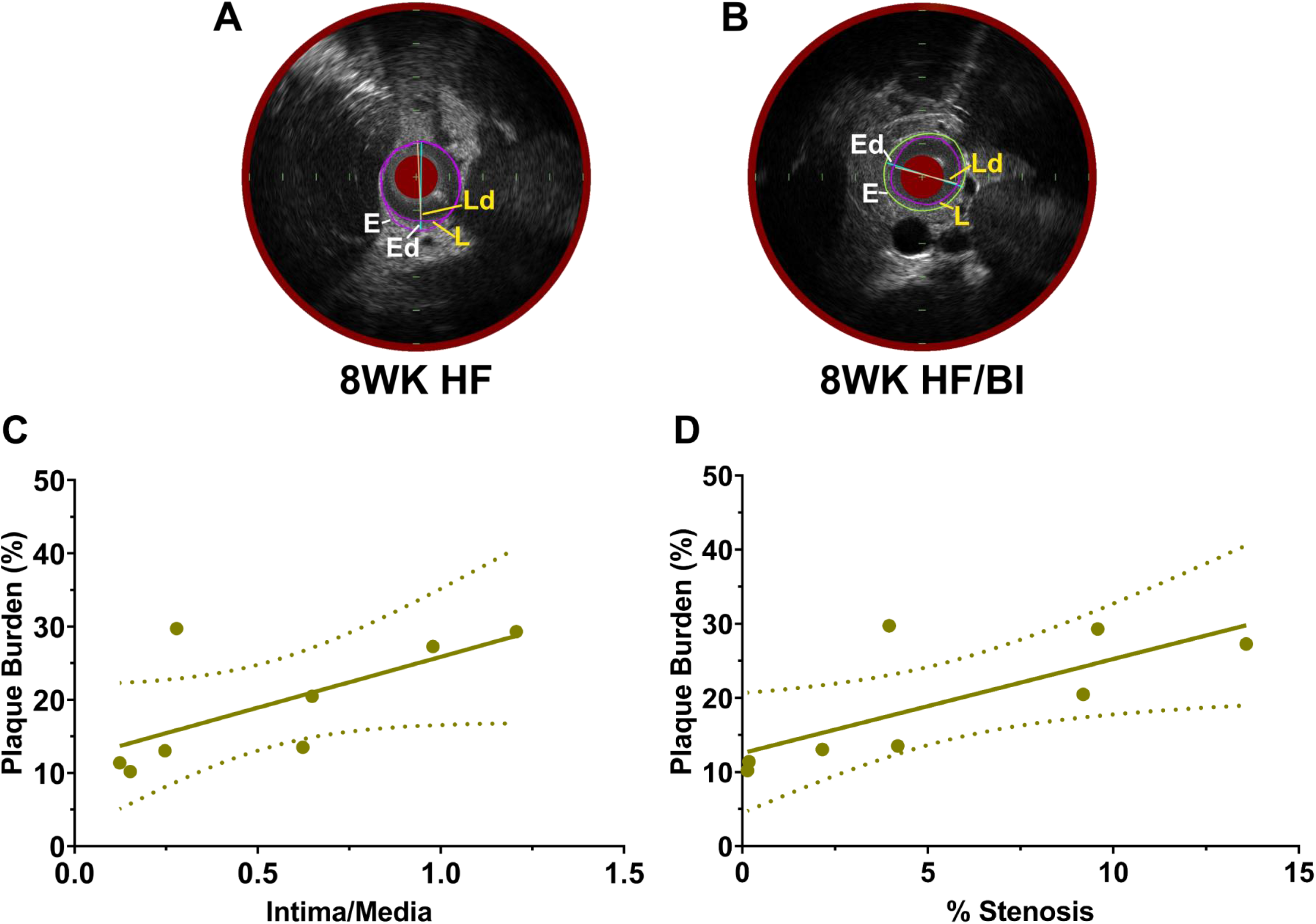
Plaque Burden Quantitation by IVUS. (A, B) Representative IVUS images. Plaque burden, defined as 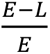, was calculated from IVUS images by comparing the areas enclosed by the external elastic lamina, *E*, and the lumen, *L*. Ed and Ld represent the maximum diameters of the areas enclosed by the external elastic lamina and the lumen, respectively. Plaque burden correlated significantly with both (C) intima/media ratio (r=0.939, *P*<0.001) and (D) %stenosis (r=0.892, *P*=0.001). Pearson correlation coefficients were calculated for comparison of IVUS-derived plaque burden against histological parameters. HF: high-fat diet. HF/BI: high-fat diet & balloon injury. IVUS: intravascular ultrasound.

**Figure 5.**
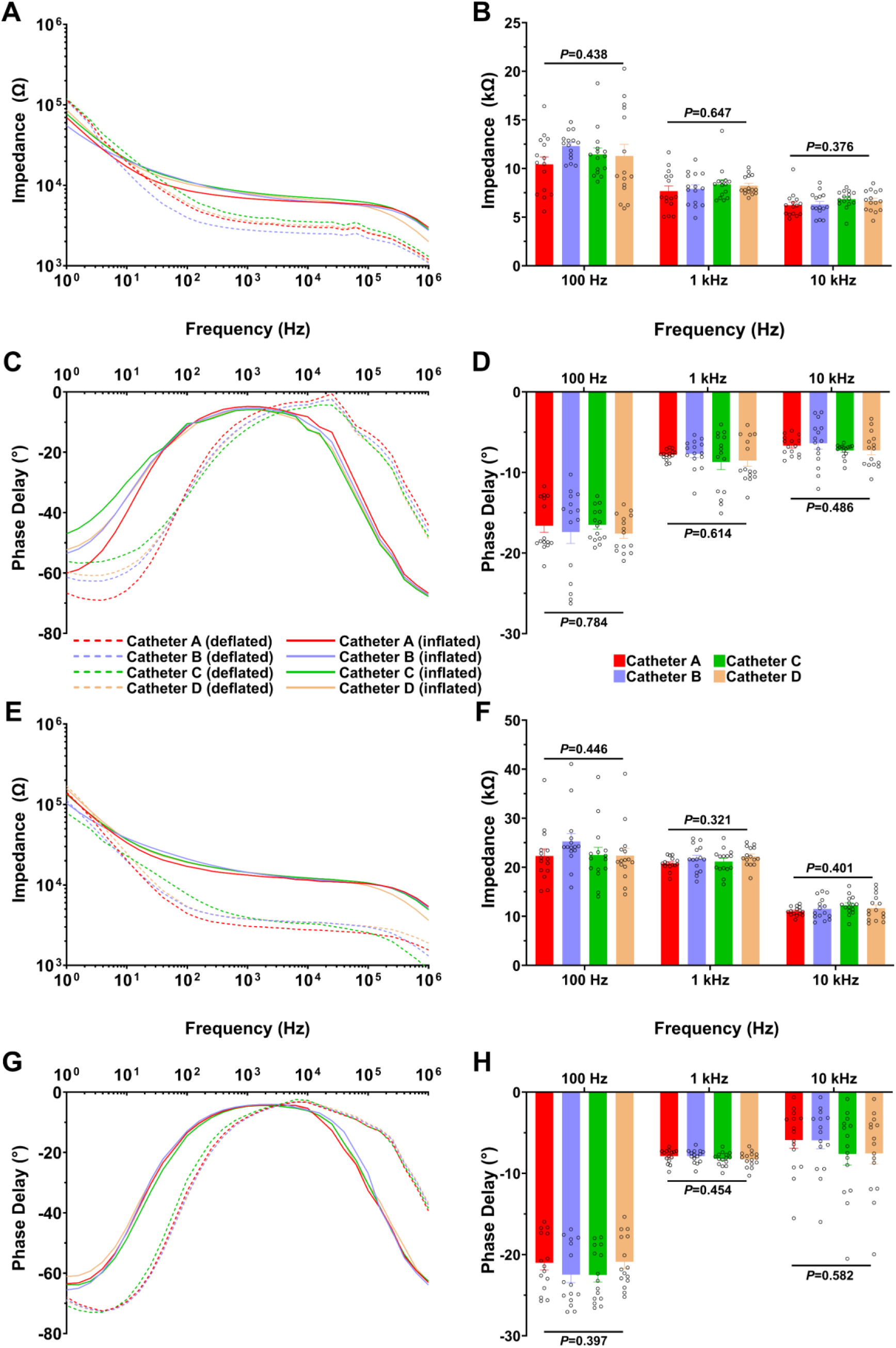
Assessment of EIS Measurement Stability among Multiple Catheters. The stability of (A-B, E-F) impedance and (C-D, G-H) phase delay measurements between the different catheters that were utilized for *in vivo* interrogation was assessed by separately measuring the *ex vivo* impedance and phase delay profiles of a control (A-D) and an 8WK HF (E-H) abdominal aorta segment in 70 mM NaCl solution. (A, C, E, G) Measurements were performed with the sensor balloon inflated (solid lines) and deflated (dashed lines). Each color represents a different EIS catheter used for *in vivo* measurements, which are named to distinguish one from another but are otherwise identical. (B, D, F, H) One-way ANOVA with post-hoc Tukey’s tests were performed at 100 Hz, 1 kHz, and 10 kHz. Neither the impedance nor the phase delay displayed significant differences amongst all catheters at all frequencies tested. EIS: electrochemical impedance spectroscopy. HF: high-fat diet.

**Figure 6.**
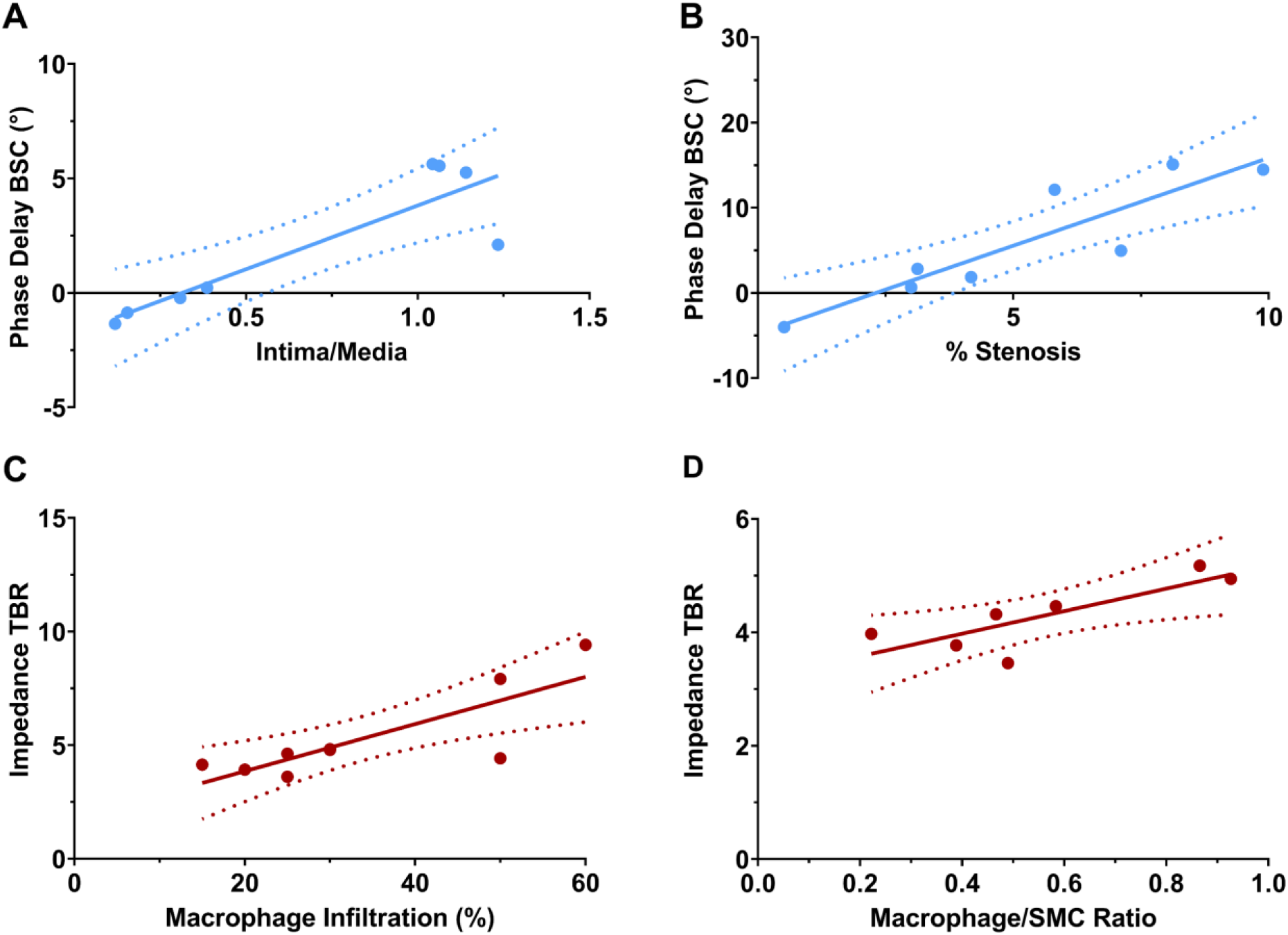
Detection of Plaque Morphology and Composition by EIS. The impedance and phase delay profiles of rabbit abdominal aortas were analyzed via EIS *in vivo* across the frequencies 1 Hz–1 MHz. (A, B) Phase delay BSC demonstrated significant correlation with intima/media ratio (0.04–2.5 kHz) and %stenosis (0.04–1.6 kHz). Correlation with intima/media ratio peaked at 1 kHz (r=0.883, *P*=0.004) and with %stenosis at 0.25 kHz (r=0.901, *P*=0.002). (C, D) The correlation of impedance TBR with macrophage infiltration (0.4–25 kHz) peaked at 10 kHz (r=0.813, *P*=0.008) and with macrophage/SMC ratio at 25 kHz (r=0.813, *P*=0.026). Pearson correlation coefficients were calculated for comparison of EIS impedance and phase delay against histological parameters. BSC: background subtraction correction. EIS: electrochemical impedance spectroscopy. SMC: smooth muscle cell. TBR: target to background ratio.

**Figure 7.**
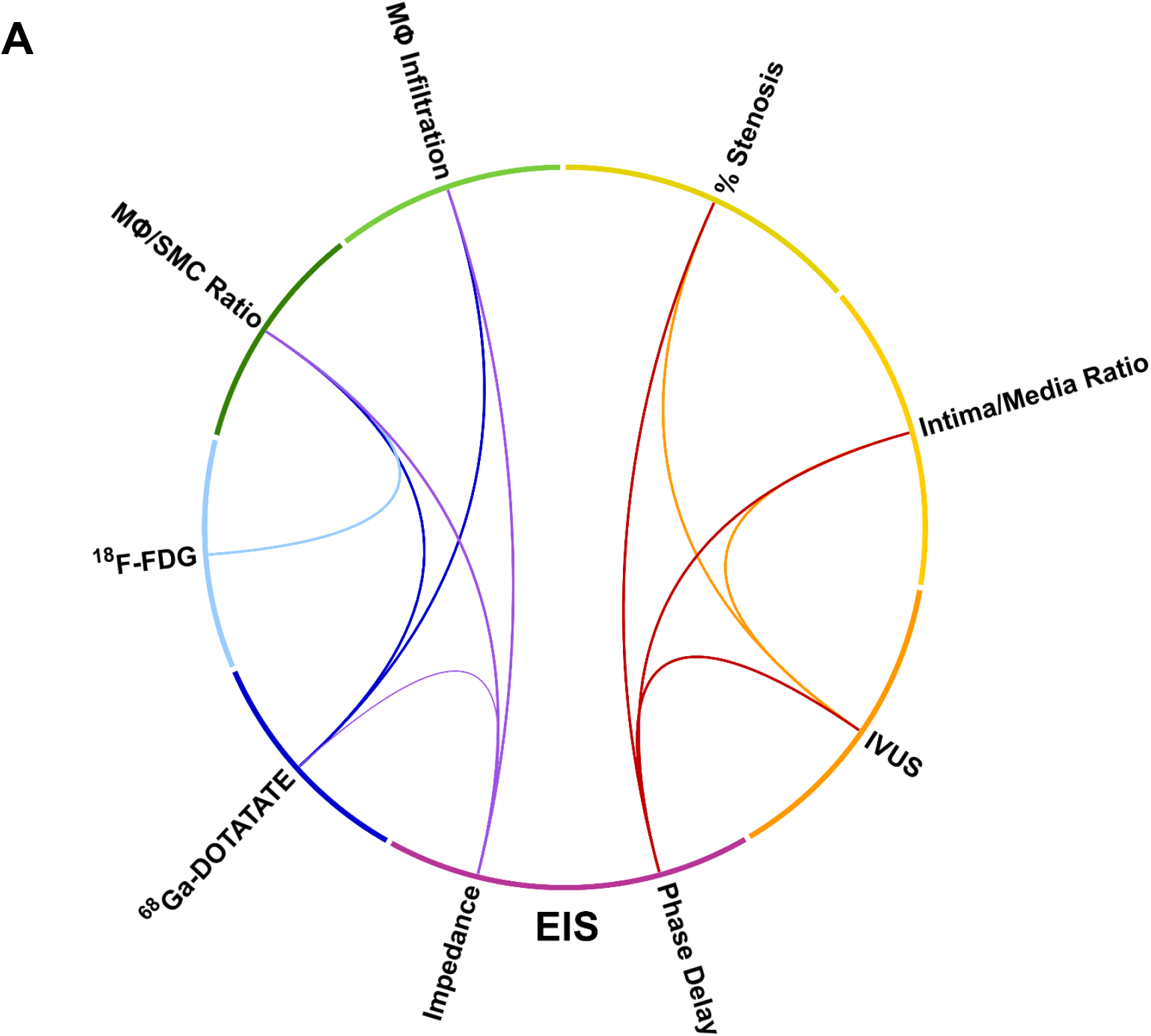
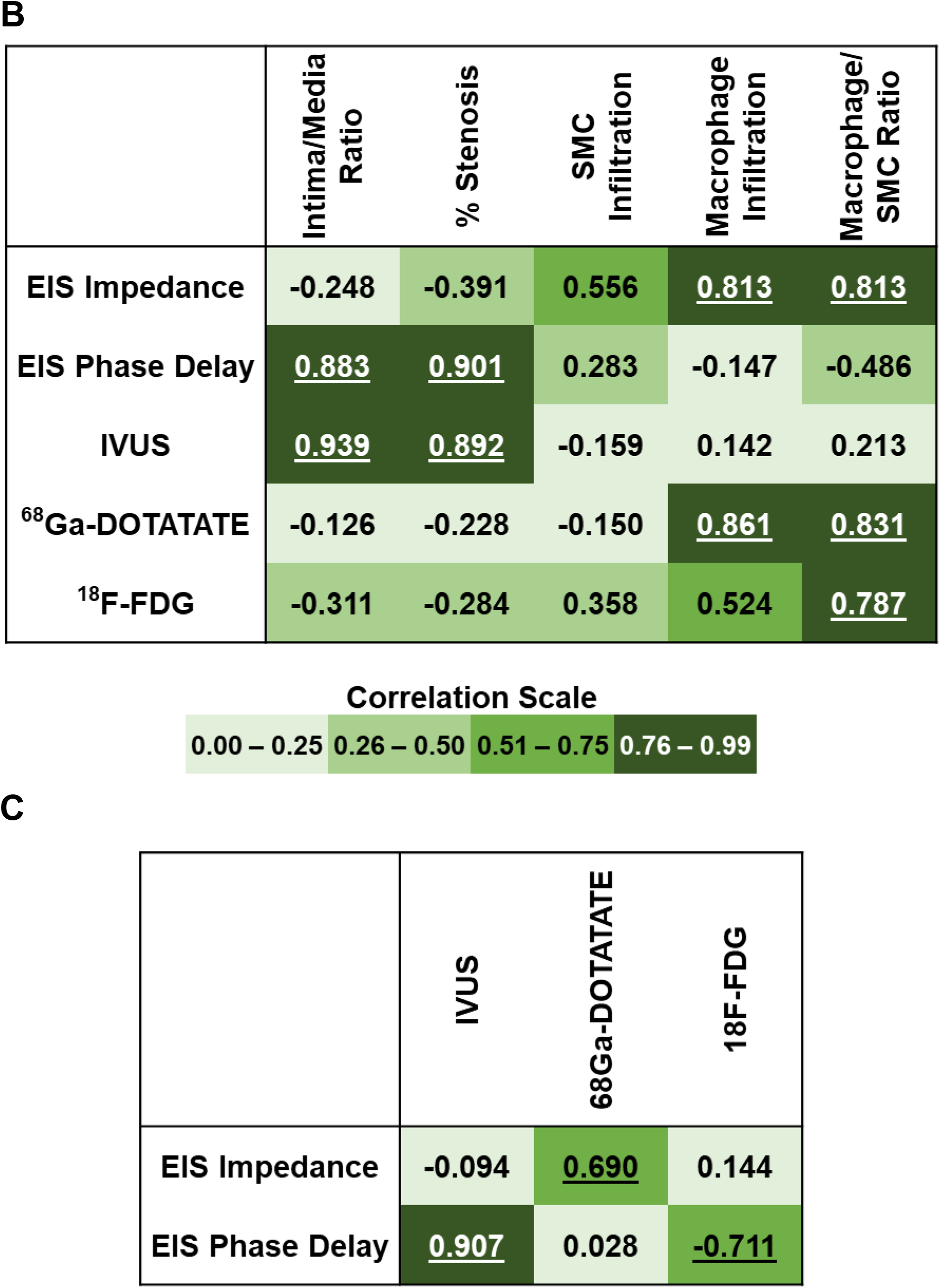
Correlation of EIS, PET, and IVUS with Histological Plaque Parameters. (A) The thickness of each line represents the magnitude of correlation, i.e., thicker lines indicate a stronger correlation. Only statistically significant correlations are shown. (B) Companion table to Panel (A). Correlation strengths are categorized into the following groups: 0.00-0.25, 0.26-0.50, 0.51-0.75, and 0.76-0.99, with corresponding color. Statistically significant correlations are underlined. (C) Additional correlations of EIS with IVUS and PET radiotracers are presented for completeness. Our intended use of EIS is the detection of metabolic activity via impedance and morphological features via phase delay. EIS: electrochemical impedance spectroscopy. ^18^F-FDG: ^18^F-fluorodeoxyglucose. ^68^Ga-DOTATATE: ^68^Ga-tetraazacyclododecanetetraacetic acid-DPhe1-Tyr3-octreotate. IVUS: intravascular ultrasound. MΦ: macrophage. SMC: smooth muscle cell.

### Blood Work

Blood samples for total cholesterol, low-(LDL), and high-(HDL) density lipoprotein cholesterol, triglycerides, and C-reactive protein (CRP) (VRL Diagnostics) were collected prior to high-fat diet initiation and 24 h prior to harvesting following overnight fast.

### Balloon Injury

Rabbits from the 4WK HF/BI (n=4) and 8WK HF/BI (n=4) groups were subjected to endothelial injury of the infrarenal abdominal aorta via balloon denudation. General anesthesia was induced via administration of ketamine (10 mg/kg) and dexmedetomidine (10 mcg/kg) intravenously using an ear catheter. Under general anesthesia, rabbits were placed on a mechanical ventilator via endotracheal intubation to deliver isoflurane (2-3.5%) for maintenance of anesthesia throughout the duration of the procedure. A cutdown was performed in the right inguinal area to expose the right femoral artery. A 5-French vascular sheath (Terumo) was inserted into the right femoral artery and a 3-French thru-lumen embolectomy balloon catheter (Edwards Lifesciences) was advanced under fluoroscopic guidance (Siemens Artis Zeego with robotic arm) and iodine contrast (Ultravist 300 mg/mL, McKesson) injection through the abdominal aorta to 2 cm caudal from the takeoff of the left renal artery (**Figure 2**). The balloon catheter was inflated and three back-and-forth pullbacks over a 2cm region performed. The sheath was removed, femoral artery ligated, and surgical site closed in layers with absorbable sutures.

### Micro-PET/CT Imaging

All animals were imaged 72 h (^68^Ga-DOTATATE), 48 h (^18^F-NaF), and 24 h (^18^F-FDG) prior to harvesting. Animals were fasted overnight prior to ^18^F-FDG imaging. Animals were injected with either 37 MBq (^68^Ga-DOTATATE, ^18^F-NaF) or 111 MBq (^18^F-FDG) of radiotracer diluted in 0.5-1 mL sterile saline solution (0.9% w/v NaCl) via the marginal ear vein. After injection, 1 h, 1.5 h, or 3 h was allowed for uptake of ^68^Ga-DOTATATE, ^18^F-NaF, or ^18^F-FDG, respectively. Following the uptake period, animals were anesthetized using the aforementioned procedure and maintained on anesthesia using isoflurane delivered through a nose cone, placed prone on the scanner bed, and positioned such that the scanner field of view (12 cm width) was centered 2 cm caudal to the left renal takeoff. Micro-PET (350-650 keV, 7-minute scan time, 0.5435 mm voxel size) and low-attenuation CT (80 kVp, 150 μA, 720 projections, 1-minute scan time) images were acquired on a GNEXT micro-PET/CT scanner (Sofie Biosciences). After micro-PET/CT image acquisition, 5 mL of Ultravist 300 was injected via the marginal ear vein and contrast CT image acquisition initiated. An additional 5 mL of Ultravist 300 followed by 5 mL sterile saline and 1 mL 0.1% heparin flush were injected over the first 30 seconds of the contrast CT scan. All fluids were kept on a heating pad prior to injection. The total contrast CT scan time was 90 seconds. Micro-PET/CT and contrast CT scans were repeated for the chest area using the aforementioned procedures.

Micro-PET images were reconstructed using a 3D-ordered subset expectation maximization algorithm (24 subsets and 3 iterations), with random, attenuation, and decay correction. The CT images were reconstructed using a Modified Feldkamp Algorithm. PET and CT images were interpolated using trilinear interpolation (AMIDE). The low-attenuation CT and contrast CT images were aligned using bone structures as reference. Once the datasets were aligned, a cylindrical region-of-interest (ROI) of 3 mm radius and 20 mm length was placed on the abdominal aorta 2-4 cm caudal to the left renal takeoff. To correct for background blood pool activity, a spherical ROI with radius of 2 mm was placed in the right atrium. The %ID/cc were obtained.

### IVUS

Prior to EIS measurements, an IVUS catheter (Makoto Intravascular Imaging System, Infraredx) was advanced through the sheath under fluoroscopic guidance to 2 cm caudal to the left renal artery takeoff. A 2 cm region was interrogated using a pullback speed of 0.5 mm/s. IVUS images were analyzed for plaque burden, defined as 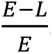, where *E* is the area enclosed by the external elastic membrane and *L* is the area of the lumen.

### EIS Measurements

EIS was performed *in vivo*, under general anesthesia using the aforementioned procedures and immediately prior to animal euthanasia. Following placement of a 5-French sheath via femoral artery, an EIS sensor catheter was inserted and advanced into the abdominal aorta under fluoroscopic guidance to 2 cm caudal of the left renal artery takeoff (**Figure 2D**). The balloon was inflated until contact was made with the endoluminal surface. Two replicates of all 15 pair-wise permutations of EIS measurements were performed from 1 Hz─1 MHz using AC signals with peak-to-peak voltages of 50 mV and five data points per decade (**Figure 2E)**. Impedance (Ω) and phase delay (°) were quantified. Prior to inflating the balloon, a set of measurements were also obtained with the balloon deflated to verify specificity of EIS signals. The diagnostic performance of impedance and phase delay raw values, target-to-background ratios (TBR: inflated ÷ deflated), and background subtraction correction (BSC: inflated − deflated) was determined. Separately, *ex vivo* EIS measurements of abdominal aortas from a control rabbit and an 8WK HF rabbit were performed with all catheters that were utilized for the *in vivo* EIS measurements to assess the stability of EIS measurements across different catheter units.

### Histology

The region of the abdominal aorta 2-4 cm caudal to the left renal artery takeoff was collected following IVUS and EIS interrogation. Tissues were washed in phosphate-buffered saline (PBS), fixed in 10% formalin, and stored in 70% ethanol. Samples were then embedded in paraffin, sectioned, and stained with hematoxylin and eosin, Masson’s trichrome, and von Kossa (Statlab) staining methods. Immunohistochemistry was performed to detect vascular SMC (α-actin, Agilent Dako). Macrophage infiltration was evaluated visually by an experienced pathologist. Atherosclerotic plaques were analyzed for intima/media thickness ratio, percent stenosis (%stenosis), smooth muscle cell (SMC) infiltration, macrophage infiltration, and macrophage/SMC ratio. %stenosis was defined as the ratio of the endoluminal surface circumference to the internal elastic lamina circumference. The thicknesses of the intimal and medial layers were measured at four locations equidistant from each other along the internal and external elastic laminas. The intima/media thickness ratio was calculated by dividing the maximum intimal thickness by the maximum medial thickness. % SMC infiltration was obtained by calculating the percent intimal area that displayed positive α-actin staining. All analyses were conducted blinded to experimental conditions.

### Statistics

Results are presented as mean ± standard error or Pearson correlations. Two-tailed unpaired Student’s t-tests were performed to assess differences amongst groups in serum biomarker levels. One-way ANOVA with post-hoc Tukey’s test was performed to evaluate PET radiotracer activity, IVUS-derived plaque burden across groups, and consistency of EIS measurements amongst the different catheters utilized. The Shapiro-Wilk test was performed to assess normality of data sets. The Pearson correlation coefficient was used for comparison of IVUS-derived plaque burden, micro-PET/CT results, impedance, and phase delay against histological parameters. Prism version 9 (GraphPad) was used for statistical analyses. A P-value <0.05 was considered significant.

## ACKNOWLEDGMENTS

We thank Stephanie Grainger, Stephen Muse, and Paul Poronto, all from Makoto Intravascular Imaging System, Infraredx, for the loan and use of the IVUS animal imaging system.

## SOURCES OF FUNDING

Dr. Packard is supported by VA Merit BX004558, NIH R56HL158569, NIH NCATS UCLA CTSI UL1TR001881, and UCLA Cardiovascular Discovery Fund/Lauren B. Leichtman and Arthur E. Levine Investigator Award. The GNEXT microPET/CT scanner (Sofie Biosciences) was funded by an NIH Shared Instrumentation for Animal Research (SIFAR) Grant (1S10OD026917-01A1) and supported by the Cancer Center Support Grant (2P30CA016042-44).

## DISCLOSURES

None.

## Non-standard Abbreviations and Acronyms

AC: alternating current
ACS: acute coronary syndrome
BSC: background subtraction correction
CAD: coronary artery disease
CRP: C-reactive protein
CT: computed tomography
CTA: computed tomography angiography
EIS: electrochemical impedance spectroscopy
FFR: fractional flow reserve
FFR-CT: fractional flow reserve by computed tomography
^18^F-FDG: ^18^F-fluorodeoxyglucose
^18^F-NaF: ^18^F-sodium fluoride
^68^Ga-DOTATATE: ^68^Ga-tetraazacyclododecanetetraacetic acid-DPhe1-Tyr3-octreotate
HDL: high-density lipoprotein
HF: high-fat diet
HF/BI: high-fat diet & balloon injury
%ID/cc: % injected dose per cubic centimeter
IVUS: intravascular ultrasound
LDL: low-density lipoprotein
NIRF: near-infrared fluorescence
NZW: New Zealand White
OCT: optical coherence tomography
PBS: phosphate-buffered saline
PET: positron emission tomography
ROI: region-of-interest
SMC: smooth muscle cell
SSTR2: somatostatin receptor 2
TBR: target to background ratio

## Notes

### Competing Interest Statement

The authors have declared no competing interest.

