## Supplemental Material for "Flexible 3-D Electrochemical Impedance Spectroscopy Sensors Incorporating Phase Delay for Comprehensive Characterization of Atherosclerosis"

**Supplemental Figure 1. Serum Lipid and CRP Levels at Baseline and Harvest**. Fasting serum levels of (A) total cholesterol (B) LDL cholesterol (C) HDL cholesterol (D) triglycerides and (E) CRP were assessed prior to initiation of high-fat diet and at harvest. Only total cholesterol and LDL cholesterol displayed significantly different serum values between baseline and harvest for all four experimental groups. Both serum HDL cholesterol and triglyceride levels were significantly different between the two balloon injury groups at harvest. Differences in serum marker levels between groups were evaluated via two-tailed unpaired Student's t-tests. **P*<0.05, ***P*<0.01 for within group comparisons; #*P*<0.05 for within timepoint comparisons. CRP: C-reactive protein. HDL: high-density lipoprotein. HF: high-fat diet. HF/BI: high-fat diet & balloon injury. LDL: low-density lipoprotein.

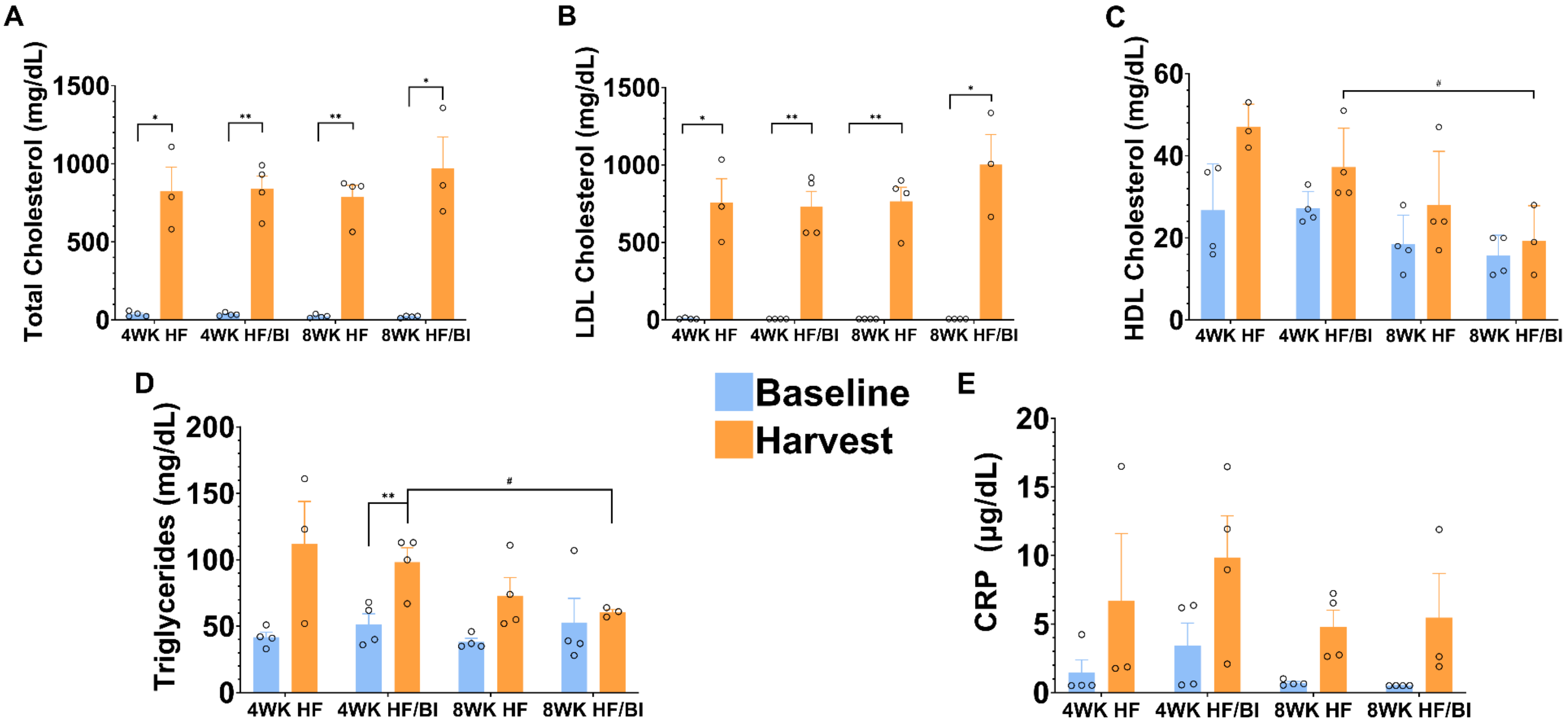

**Supplemental Figure 2. Histological Sections of Abdominal Aorta.** Representative samples of abdominal aorta stained with (A, B) H&E, (C, D) Masson's Trichrome, (E, F) von Kossa, and (G, H) anti-smooth muscle α-actin. The neointimal layer (black bars) is bound by the internal elastic lamina (green arrows). Calcification is marked by an asterisk. SMC infiltration was quantified by comparing the area of brown staining to the total area of the neointima. Orange arrows indicate the external elastic lamina. Images from the top row were taken at 6X magnification (scale bar: 400 μm), and those from the bottom row at 30X magnification (scale bar: 70 μm). H&E: hematoxylin & eosin. SMA: smooth muscle α-actin. SMC: smooth muscle cell.

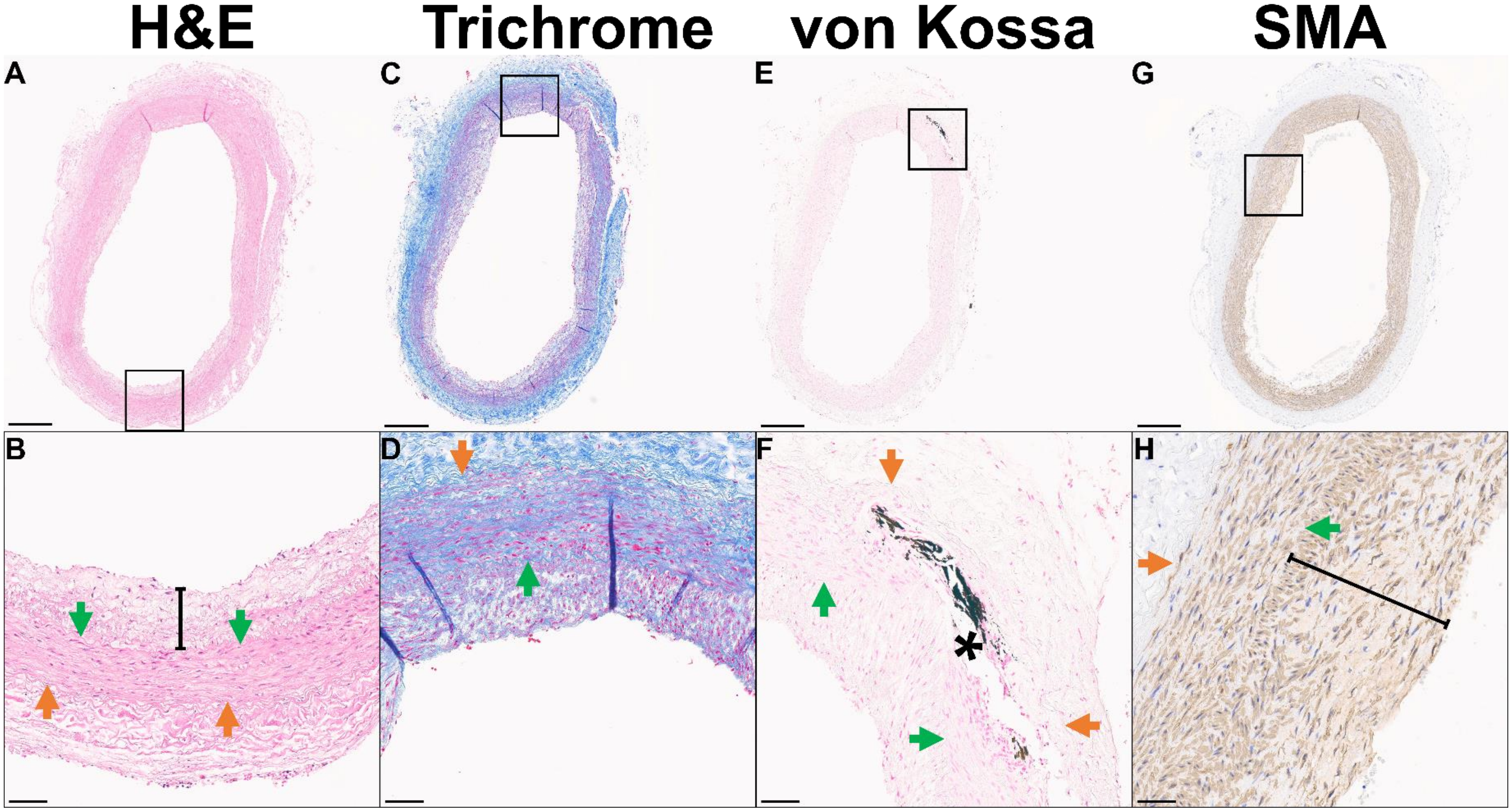

**Supplemental Figure 3. Histological Analysis of Rabbit Abdominal Aortas.** Histological sections of rabbit abdominal aortas were analyzed for the (A) intima/media thickness ratio, (B) % stenosis, (C) % macrophage infiltration, (D) % SMC infiltration, and (E) macrophage/SMC infiltration ratio. (A, B) Other than the 4WK HF group and one 8WK HF rabbit, intimal thickening was present in the abdominal aortas of all rabbits. Whereas the 8WK HF group displayed minute, focal intimal thickening (27.6 ± 6.7μm), substantial, circumferential intimal thickening was present in both the 4WK HF/BI (125.5 ± 67.8μm) and 8WK HF/BI (200.6 ± 81.2μm) groups. Similarly, intima/media ratio and %stenosis, absent in the 4WK HF group, were lowest in the 8WK HF group (0.14 ± 0.03, 1.76% ± 1.76%), followed by the 4WK HF/BI group (0.64 ± 0.51, 4.80% ± 2.07%), and most significant in the 8WK HF/BI group (1.08 ± 0.05, 7.94% ± 2.05%). (D, E) In the HF only groups, there was no macrophage or intimal SMC infiltration at 4WK, and these were present only in a subset at 8WK (n=2 for macrophage, 35.00% ± 21.21%). On the other hand, all rabbits from the 4WK HF/BI and 8WK HF/BI groups exhibited macrophage (37.50% ± 21.02%, and 28.33% ± 2.89%, respectively) and SMC (63.61% ± 4.15%, and 55.60% ± 7.56%, respectively) accumulation in the intimal layer. HF: high-fat diet. HF/BI: high-fat diet & balloon injury. MΦ: macrophage. SMC: smooth muscle cell.

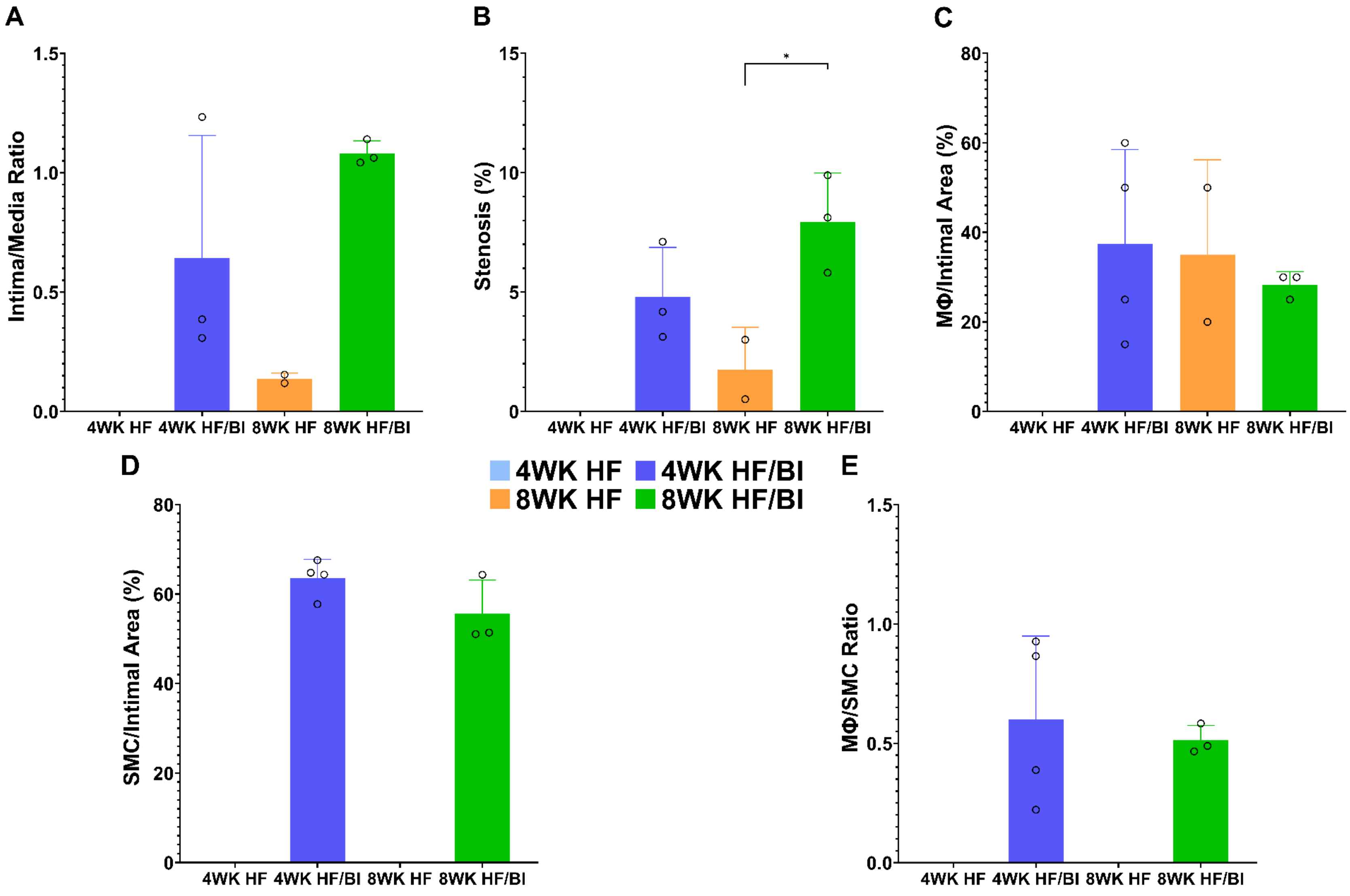

A
Intima/Media Ratio
1.5
1.0
0.5
0.0
4WK HF 4WK HF/BI 8WK HF 8WK HF/BI
B
Stenosis (%)
15
10
5
0
*
4WK HF 4WK HF/BI 8WK HF 8WK HF/BI
C
MΦ/Intimal Area (%)
80
60
40
20
0
4WK HF 4WK HF/BI 8WK HF 8WK HF/BI
D
SMC/Intimal Area (%)
80
60
40
20
0
4WK HF 4WK HF/BI 8WK HF 8WK HF/BI
4WK HF
4WK HF/BI
8WK HF
8WK HF/BI
E
MΦ/SMC Ratio
1.5
1.0
0.5
0.0
4WK HF 4WK HF/BI 8WK HF 8WK HF/BI

**Supplemental Figure 4. Atherosclerosis Characterization by PET Radiotracers**. (A) $^{68}$Ga-DOTATATE uptake peaked in the 8WK HF group. A significant increase was observed from the 4WK HF group to the 8WK HF group (0.023 ± 0.002 vs 0.038 ± 0.004, *P*=0.020), while a significant decrease was observed from the 8WK HF group and to 8WK HF/BI groups (0.038 ± 0.004 vs 0.024 ± 0.002, *P*=0.032). (B) $^{18}$F-NaF uptake was highest in the 4WK HF/BI and the 8WK HF/BI groups. $^{18}$F-NaF activity above background levels was not observed in the 4WK HF and 8WK HF groups. There was a significant increase between the 8WK HF and 8WK HF/BI groups (1.081 ± 0.352 vs 2.66 ± 0.534, *P*=0.049). (C) $^{18}$F-FDG uptake significantly increased from the 4WK HF group to the 8WK HF group, where it peaked, (0.848 ± 0.076 vs 1.364 ± 0.030, *P*=0.003) and then decreased from the 8WK HF group to the 8WK HF/BI group (1.364 ± 0.030 vs 0.773 ± 0.118, *P*=0.008). Differences in PET radiotracer activity between groups were evaluated via one-way ANOVA with post-hoc Tukey's test. **P*<0.05. ***P*<0.01. $^{18}$F-FDG: $^{18}$F-fluorodeoxyglucose. $^{68}$Ga-DOTATATE: $^{68}$Ga-tetraazacyclododecanetetraacetic acid-DPhe1-Tyr3-octreotate. $^{18}$F-NaF: $^{18}$F-sodium fluoride. HF = high-fat diet. HF/BI = high-fat diet & balloon injury. TBR = target-to-background ratio.

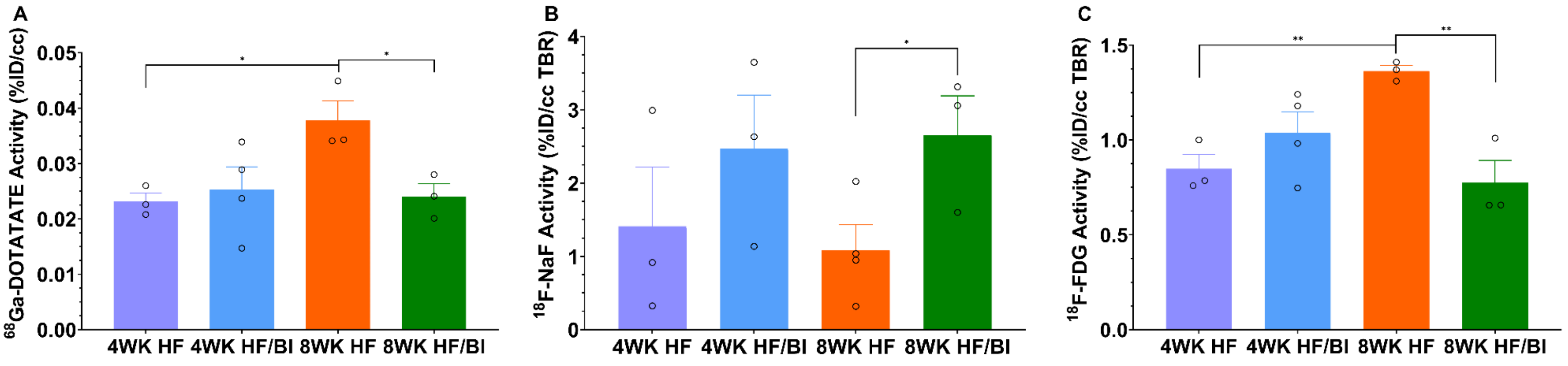

**Supplemental Figure 5. Atherosclerotic Characterization by $^{18}$F-NaF and Comparison to EIS.** (A) Representative images of the abdominal aorta obtained using $^{18}$F-NaF obtained to evaluate degree of microcalcification. Activities of $^{18}$F-NaF correlated significantly with histological plaque parameters of interest. Mean %ID/cc TBR of $^{18}$F-NaF strongly correlated with (B) calcified area (r=0.998) and (C) SMC infiltration (r=-0.842, P=0.018). Pearson correlation coefficients were calculated for comparison of $^{18}$F-NaF activity against histological parameters. (D) Correlations of $^{18}$F-NaF radiotracer activity with all histological parameters evaluated as well as EIS impedance and phase delay measurements. Correlation strengths are categorized into the following groups: 0.00-0.25, 0.26-0.50, 0.51-0.75, and 0.76-0.99, with corresponding color. Statistically significant correlations are underlined. All CT images and all PET images were obtained using the same scale. EIS: electrochemical impedance spectroscopy. $^{18}$F-NaF: $^{18}$F-sodium fluoride. HF: high-fat diet. HF/BI: high-fat diet & balloon injury. %ID/cc: % injected dose per cubic centimeter. SMC: smooth muscle cell. TBR: target-to-background ratio.

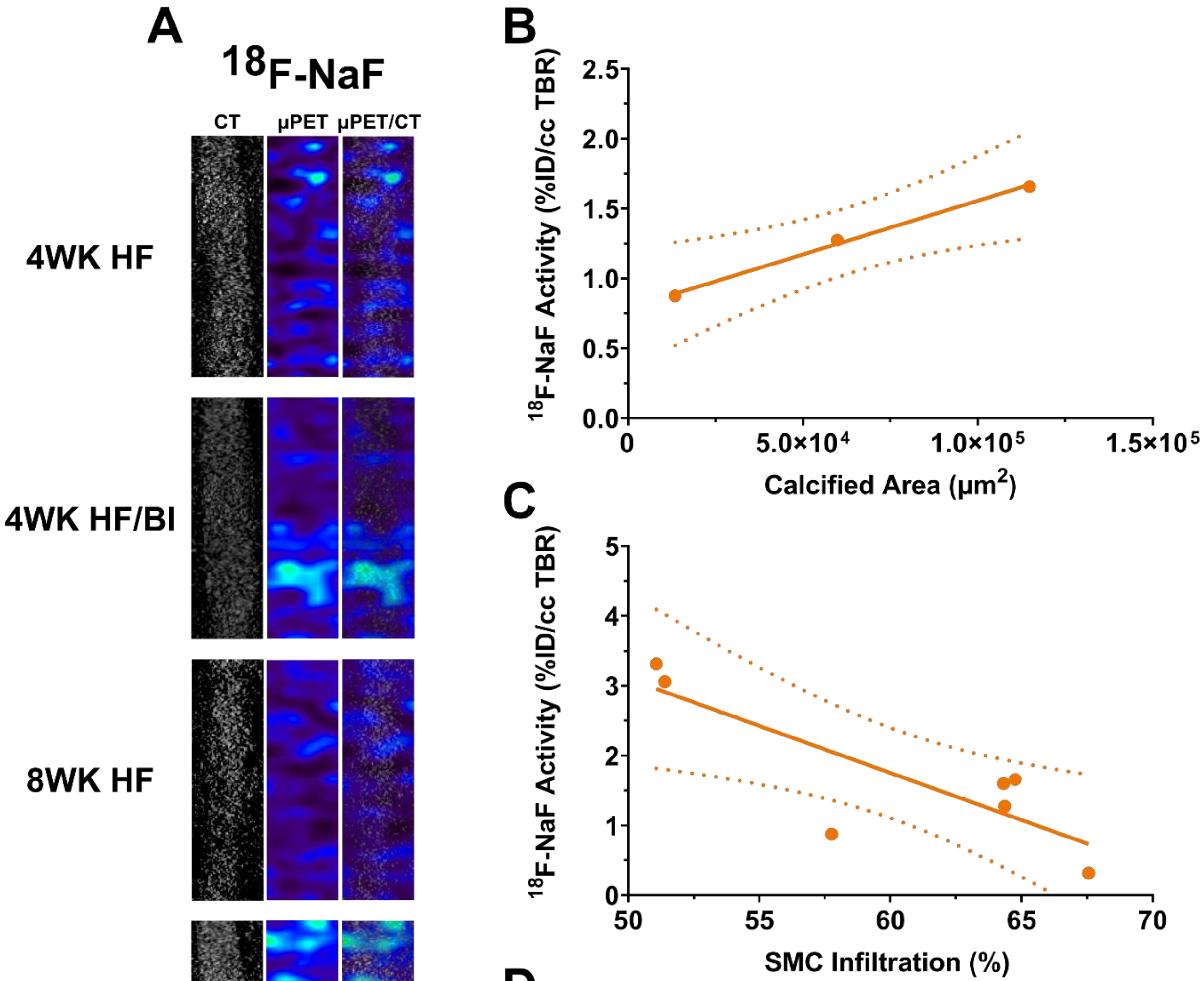


D

| | Intima/Media Ratio | % Stenosis | Calcified Area | SMC Infiltration | Macrophage Infiltration | Macrophage/SMC Ratio | EIS Impedance | EIS Phase Delay |
|---|---|---|---|---|---|---|---|---|
| $^{18}$F-NaF | 0.277 | 0.427 | 0.998 | -0.842 | -0.008 | 0.150 | -0.557 | 0.196 |

| Correlation Scale |
|---|
| 0.00 – 0.25 |
| 0.26 – 0.50 |
| 0.51 – 0.75 |
| 0.76 – 0.99 |

**Supplemental Figure 6. IVUS-derived Plaque Burden.** Plaque burden was defined as $\frac{E-L}{E}$, where *E* and *L* represent the areas enclosed by the external elastic lamina and the lumen, respectively. Rabbits from the 4WK HF group displayed no luminal narrowing. Rabbits from the 8WK HF/BI group exhibited the highest degree of plaque burden. Comparisons between timepoints (4 vs. 8 weeks) as well as experimental conditions (high-fat only vs high-fat & balloon injury) yielded statistically significant differences for all comparisons. One-way ANOVA with post-hoc Tukey's test was used to evaluate IVUS-derived plaque burden between experimental groups. **P*<0.05; ***P*<0.01; ****P*<0.001; *****P*<0.0001. HF: high-fat diet. HF/BI: high-fat diet & balloon injury. IVUS: intravascular ultrasound.

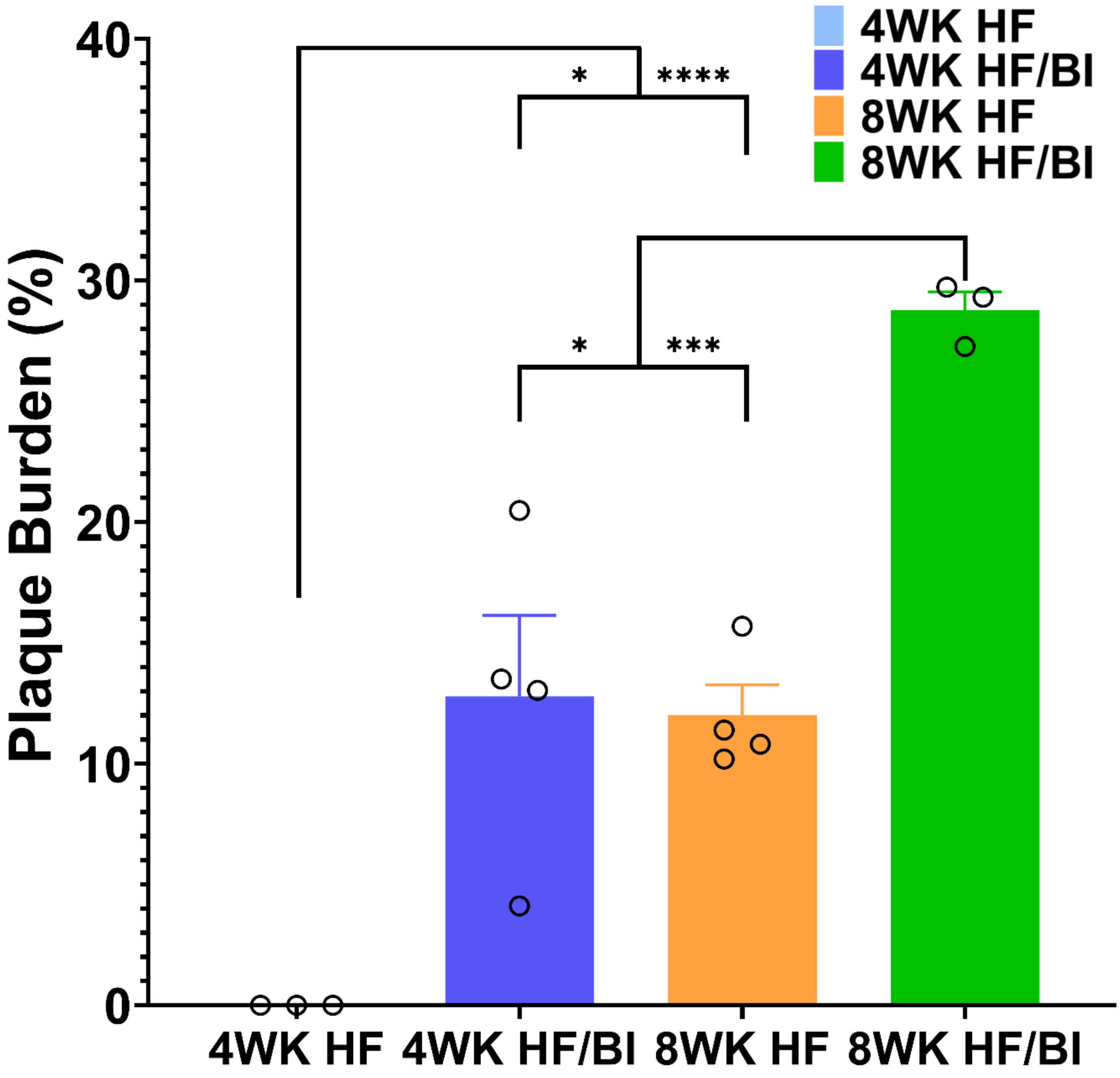

**Supplemental Figure 7. Impedance and Phase Delay Characterization of Plaques.** Impedance and phase delay profiles from representative samples of each experimental group. EIS measurements were performed *in vivo* with the sensor balloon inflated (solid lines) and deflated (dashed lines). Each color represents a different EIS sensor permutation. EIS: electrochemical impedance spectroscopy. HF: high-fat diet. HF/BI: high-fat diet & balloon injury.

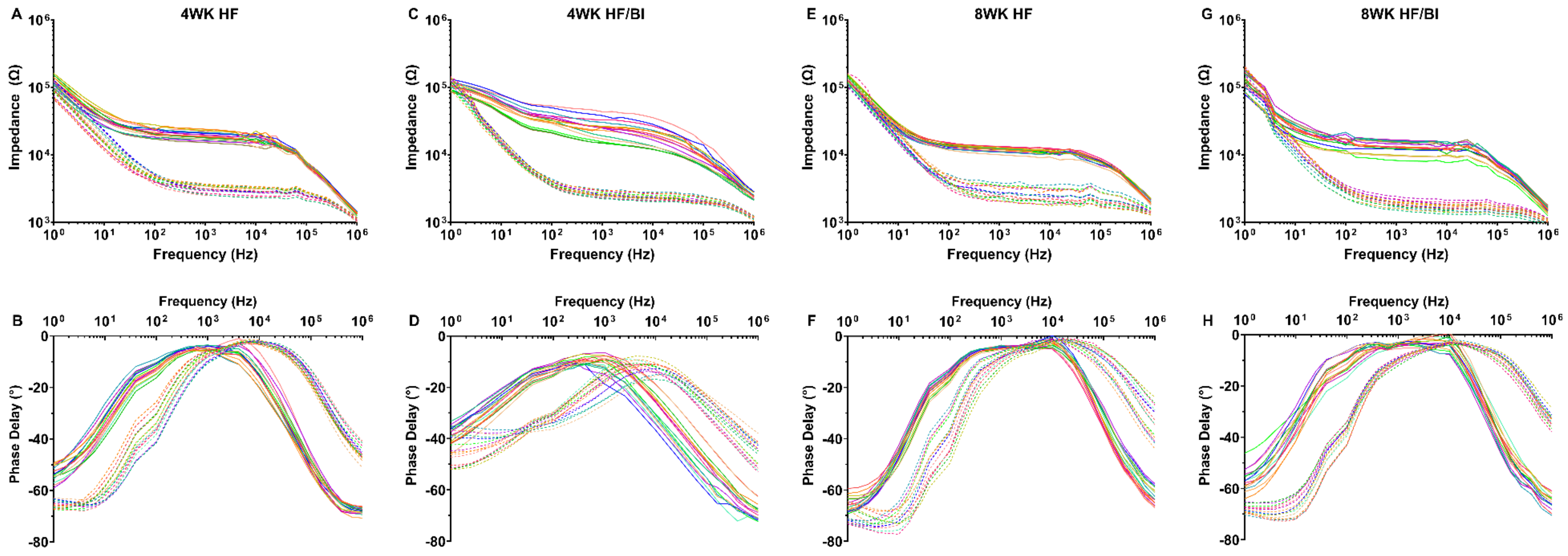

**Supplemental Table 1. Serum Biomarkers at Baseline and Harvest.** Companion Table to Supplemental Figure 1. Total cholesterol and LDL cholesterol levels significantly increased from baseline to harvest in all experimental groups. Serum triglyceride levels significantly increased within the 4WK HF/BI group. Values are presented as Mean ± SEM. *P*-values were derived from two-tailed unpaired Student's t-test between timepoints within the same experimental group. Significant *P*-values are bolded. CRP: C-reactive protein. HDL: high-density lipoprotein. HF: high-fat diet. HF/BI: high-fat diet & balloon injury. LDL: low-density lipoprotein.

| Parameter | Group | Baseline | Harvest | *P*-value |
|---|---|---|---|---|
| **Total Cholesterol (mg/dL)** | **4WK HF** | 37.25 ± 8.23 | 826.00 ± 153.60 | **0.032** |
| | **4WK HF/BI** | 36.25 ± 5.15 | 839.00 ± 82.33 | **0.003** |
| | **8WK HF** | 24.00 ± 5.57 | 787.50 ± 74.64 | **0.002** |
| | **8WK HF/BI** | 21.25 ± 3.30 | 972.00 ± 198.86 | **0.040** |
| **LDL Cholesterol (mg/dL)** | **4WK HF** | 8.25 ± 2.25 | 756.67 ± 154.07 | **0.039** |
| | **4WK HF/BI** | 6.00 ± 0.00 | 732.00 ± 97.88 | **0.005** |
| | **8WK HF** | 6.00 ± 0.00 | 765.00 ± 91.59 | **0.004** |
| | **8WK HF/BI** | 6.00 ± 0.00 | 1002.67 ± 193.71 | **0.036** |
| **HDL Cholesterol (mg/dL)** | **4WK HF** | 26.75 ± 5.65 | 47.00 ± 3.21 | 0.221 |
| | **4WK HF/BI** | 27.25 ± 2.02 | 37.25 ± 4.73 | 0.197 |
| | **8WK HF** | 18.50 ± 3.52 | 28.00 ± 6.54 | 0.058 |
| | **8WK HF/BI** | 15.75 ± 2.46 | 19.33 ± 4.91 | 0.185 |
| **Triglycerides (mg/dL)** | **4WK HF** | 41.75 ± 3.68 | 112.00 ± 31.94 | 0.136 |
| | **4WK HF/BI** | 51.50 ± 7.93 | 98.25 ± 10.86 | **0.005** |
| | **8WK HF** | 38.25 ± 2.63 | 73.00 ± 13.57 | 0.103 |
| | **8WK HF/BI** | 52.75 ± 18.24 | 60.67 ± 2.03 | 0.991 |
| **CRP (μg/dL)** | **4WK HF** | 1.46 ± 0.93 | 6.72 ± 4.90 | 0.308 |
| | **4WK HF/BI** | 3.44 ± 1.64 | 9.87 ± 3.02 | 0.082 |
| | **8WK HF** | 0.71 ± 0.10 | 4.79 ± 1.22 | 0.051 |
| | **8WK HF/BI** | 0.51 ± 0.01 | 5.48 ± 3.22 | 0.262 |
